# Novel gene-intergenic fusion involving ubiquitin E3 ligase UBE3C causes distal hereditary motor neuropathy: A new mechanism for motor neuron degeneration

**DOI:** 10.1101/2022.08.16.504208

**Authors:** Anthony N. Cutrupi, Ramesh K. Narayanan, Gonzalo Perez-Siles, Bianca R. Grosz, Kaitao Lai, Alexandra Boyling, Melina Ellis, Ruby CY Lin, Brent Neumann, Di Mao, Motonari Uesugi, Garth A. Nicholson, Steve Vucic, Mario A. Saporta, Marina L. Kennerson

**Affiliations:** Northcott Neuroscience Laboratory, ANZAC Research Institute, Sydney, Australia; Faculty of Medicine and Health, University of Sydney, Sydney, NSW, Australia; Molecular Medicine Laboratory, Concord Repatriation General Hospital, Sydney, Australia; Brain and Nerve Research Centre, Concord Repatriation General Hospital, Sydney, Australia; Department of Neurology, University of Miami Miller School of Medicine, Miami, FL, 33136, USA; Institute for Integrated Cell-Material Sciences and Institute for Chemical Research, Kyoto University, Japan; Ancestry & Health Genomics Laboratory, Charles Perkins Centre, School of Medical Sciences, University of Sydney, Australia; Centre for Infectious Diseases and Microbiology, Westmead Institute for Medical Research, Sydney, NSW, Australia; Monash Biomedicine Discovery Institute and Department of Anatomy and Developmental Biology, Monash University, Melbourne, VIC 3800, Australia

**Keywords:** iPSC-derived motor neurons, distal hereditary motor neuropathy, gene-intergenic fusion, ubiquitin E3 ligase, ubiquitin proteasome system

## Abstract

Distal hereditary motor neuropathies (dHMNs) are a group of inherited diseases involving the progressive, length-dependent axonal degeneration of the lower motor neurons. There are currently 29 reported causative genes and 4 disease loci implicated in dHMN. Despite the high genetic heterogeneity, mutations in the known genes account for less than 20% of dHMN cases with the mutations identified predominantly being point mutations or indels. We have expanded the spectrum of dHMN mutations with the identification of a 1.35 Mb complex structural variation (SV) causing a form of autosomal dominant dHMN (DHMN1 OMIM %182906). Given the complex nature of SV mutations and the importance of studying pathogenic mechanisms in a neuronal setting, we generated a patient-derived DHMN1 motor neuron model harbouring the 1.35 Mb complex insertion. The DHMN1 complex insertion creates a duplicated copy of the first 10 exons of the ubiquitin-protein E3 ligase gene (*UBE3C*) and forms a novel gene-intergenic fusion sense transcript by incorporating a terminal pseudo-exon from intergenic sequence within the DHMN1 locus. The *UBE3C* intergenic fusion (*UBE3C-IF*) transcript does not undergo nonsense-mediated decay and results in a significant reduction of wild type full length UBE3C (UBE3C-WT) protein levels in DHMN1 iPSC-derived motor neurons. An engineered transgenic *C. elegans* model expressing the UBE3C-IF transcript in GABA-ergic motor neurons shows neuronal synaptic transmission deficits. Furthermore, the transgenic animals are susceptible to heat stress which may implicate defective protein homeostasis underlying DHMN1 pathogenesis. Identification of the novel *UBE3C-IF* gene-intergenic fusion transcript in motor neurons highlights a potential new disease mechanism underlying axonal and motor neuron degeneration. These complementary models serve as a powerful paradigm for studying the DHMN1 complex SV and an invaluable tool for defining therapeutic targets for DHMN1.

## Introduction

Distal hereditary motor neuropathies (dHMNs) are a group of clinically and genetically heterogeneous, inherited neurogenerative diseases predominantly involving the lower motor neurons of the peripheral nervous system.^1–4^ Patients present with a slowly progressive, length-dependent degeneration or ‘dying back’ of lower motor neuron axons, causing denervation of the distal limb muscles resulting in muscle atrophy, paresis and chronic disability.^1,2,5^ The dHMNs are comparatively rare to other forms of inherited peripheral neuropathy (IPN), with a prevalence of approximately 2 in every 100,000.^6^ Currently, 29 reported causative genes and 4 disease loci have been implicated in dHMN with diverse roles in motor neuron biology and function^7^ including: axonal transport^7–13^, neuronal DNA/RNA processing and transcription^14–16^, protein biosynthesis and post-translational processing^17–19^, mitochondrial function and energy production^20,21^, cell survival, apoptosis and signalling^22–24^, axonal growth and guidance^25,26^, and neuronal/axonal structural integrity and repair.^27,28^

Despite the genetic heterogeneity, mutations in known genes account for less than 20% of known dHMN cases.^23^ Furthermore, the types of mutations identified (point mutations or indels) have only been reported to affect the protein coding sequences of these genes. A large proportion of genetically undiagnosed dHMN families are likely to be caused by alternative mutational mechanisms that may occur in the non-coding region of DNA or involve structural variation (SV) rearrangements.

Structural variation is a broad term encompassing genomic rearrangements that disrupt chromosomal organization and genome architecture.^29^ There are many types of SV^30–32^ and range in size from 50 to millions of base pairs as defined by the Structural Variation Analysis Group.^33^ SV is reported to contribute to a wide array of diseases with a genetic aetiology, including sporadic developmental syndromes to Mendelian diseases.^34^ SV causing IPN is not unprecedented with over 20 cases reported to date across the spectrum of IPN, ranging from simple copy number variation to more complex chromosomal events (see Cutrupi *et al.,*^29^ for a review). The most common subtype of IPN is Charcot-Marie-Tooth 1A (CMT1A) (OMIM #118220) which is caused by a 1.5 Mb tandem duplication of chromosome 17p11.2.^35^ The duplication results in trisomy of the peripheral myelin protein gene *PMP22.*^36–39^ Similarly, atypical genomic rearrangements occurring at the CMT1A locus including two duplications upstream of *PMP22*^40,41^ also produce CMT1A. More recently, duplication of a long-range enhancer for the *PMP22* gene^42^ has been reported. Collectively, this suggests that gene dosage changes of genetic elements that control gene expression can also produce disease.

We previously reported a novel SV mutation segregating in a large Australian family (F-54) with autosomal dominant DHMN Type 1 (DHMN1, OMIM:#182960) that maps to chromosome 7q34-q36.2.^2^ Using whole genome sequencing (WGS), we identified a 1.35 Mb duplication of chromosome 7q36.3 inserted in the reverse orientation into the DHMN1 locus. The inserted sequence fragment contains four protein-coding genes (*MNX1* (also known a*s HB9*)*, NOM1, RNF32, LMBR1*) and their regulatory elements, as well as the upstream regulatory elements and first 10 exons of the ubiquitin-protein E3 ligase gene (*UBE3C*). We hypothesised the DHMN1 complex insertion may produce neuropathy through aberrant expression of gene(s) by three possible mechanisms: 1) gene dosage due to trisomy of the 1.35 Mb complex insertion or 2) position variegation of genes flanking the DHMN1 insertion breakpoints or 3) the genomic rearrangement introducing regulatory elements causing ectopic expression of flanking genes.

Investigating how the complex insertion causes the DHMN1 phenotype poses several challenges. Firstly, due to its size, current cloning techniques cannot reproduce the complex rearrangement. Secondly, the complex nature of the SV could mean that several genes show expression changes making identifying the causative gene difficult^29^ and obscuring the precise mechanism by which the SV might produce the DHMN1 phenotype. Finally, studying SV in the context of peripheral nerve degeneration has been hampered by the invasive procedures needed to examine the appropriate neuronal tissues.^29,43,44^

To address these issues, we generated an *in vitro* human spinal motor neuron (sMN) model using induced pluripotent stem cells (iPSC) reprogrammed from DHMN1 patient fibroblasts harbouring the 1.35 Mb complex insertion. We show that the genomic rearrangement results in the production of a novel gene-intergenic fusion transcript in which the *UBE3C* partial copy is transcribed from the reverse strand and incorporates a terminal pseudo-exon from sequence within the DHMN1 locus. This *UBE3C* intergenic fusion (*UBE3C-IF*) transcript is not degraded by nonsense-mediated decay, and DHMN1 spinal motor neurons (sMN) harbouring the *UBE3C-IF* transcript show significant reduction of wild-type full length UBE3C (UBE3C-WT) protein levels. Transgenic *C. elegans* expressing the *UBE3C-IF* transcript in GABA-ergic motor neurons show neuronal synaptic transmission deficits and susceptibility to heat stress which may implicate defective protein homeostasis in DHMN1 pathogenesis.

## Materials and methods

### Tissue culture and cell line maintenance

HeLa cell lines were maintained in H-DMEM consisting of Dulbecco’s Modified Eagle Medium (DMEM; Gibco, Life Technologies) supplemented with 10% (v/v) fetal bovine serum (FBS; SAFC Biosciences) and 1% (v/v) Penicillin/Streptomycin (P/S; Gibco, Life Technologies). Primary fibroblasts were maintained in F-DMEM culture medium comprising DMEM, 10% (v/v) FBS, 1% (v/v) P/S and 1% (v/v) L-glutamine (Gibco, Life technologies). All cells were maintained in 5% CO_2_, humidified air at 37°C. All research and cell culture procedures from patient skin biopsies was performed after informed consent and in accordance with relevant guidelines and regulations approved by the Sydney Local Health District Human Ethics Committee (HREC/17/CRGH/8).

### Generation, culturing, and maintenance of iPSC lines

Reprogramming of patient and unrelated, neurologically normal control fibroblasts was performed by FUJIFILM Cellular Dynamics (Wisconsin, USA) and has been previously described.^45^ In brief, DHMN1 patient fibroblasts harbouring the 1.35 Mb insertion were transfected with oriP/EBNA1(Epstein-Barr nuclear antigen-1)-based episomal vectors carrying reprogramming transgenes (*OCT4, OX2, NANOG, LIN28, c-MYC, KLF4* and *SV40LT*)^46,47^ and then seeded onto Matrigel (Corning)-coated plates in reprogramming medium. After 7 days, the medium was replaced with E8 medium (Gibco, Life Technologies) and cells were cultured for an additional 14 days. Single iPSC colonies were picked and propagated in E8 culture medium on Matrigel-coated plates. G-banded karyotyping confirmed iPSC lines were karyotypically normal (Wi-Cell; Wisconsin, USA). Pluripotency of iPSC lines was confirmed by in-house analysis of stem cell specific pluripotency genes for endogenous expression (FUJIFILM Cellular Dynamics). iPSC lines were seeded on pre-treated (0.167 mg/mL Matrigel) 6-well plates and cultured in TeSR-E8 iPSC culture medium (StemCell Technologies) according to the manufacture’s guidelines. Briefly, media was replaced daily until cells reached optimal morphology and approximately 75-80% confluence (4-5 days of culture). The cells were then passaged as aggregates (50-200 µm in diameter) using 0.5 mM EDTA and either re-seeded with split ratios (1:3 to 1:8) onto 6-well plates or cryopreserved using CryoStor CS10 Cryopreservation Reagent (Sigma). iPSC lines were maintained in 5% CO_2_, humidified air at 37°C.

### Motor neuron progenitor induction and sMN differentiation

We differentiated 3 control (C1, C2, and C3) and 3 DHMN1 patient (P1, P2, and P3) iPSC lines into sMN using previously described methods^48^ that have been adapted and modified by our laboratory to increase culturing efficacy. The protocol is based on a combination of dual SMAD inhibition, WNT activation and NOTCH inhibition to generate highly pure and homogenous populations of expandable motor neuron progenitors (MNP) that can be differentiated into mature sMN. Detailed methods are included in Supplementary Methods. Media and supplements used are summarised in Supplementary Table S3. All adherent culturing was performed in 6-well plates (Corning-Costar) pre-treated with 0.167 mg/mL Matrigel. In brief, neural induction was initiated using a chemically defined neural base medium (NbM) supplemented with 2 µM Dorsomorphin (StemCell Technologies), 3 µM CHIR990021 (Sigma) and 2 µM SB431542 (StemCell Technologies). Cells were maintained in NbM for 6 days after which they were passaged with 1 U/mL Dispase (StemCell Technologies) and cultured in NbM supplemented with 0.1 µM Retinoic Acid (RA, Sigma), 0.5 µM Smoothened Agonist (SAG, StemCell Technologies), 2 µM Dorsomorphin, 1 µM CHIR990021 and 2 µM SB431542 for a further 6 days to generate MNP. Terminal differentiation of MNP to sMN was carried out in suspension using ultra-low attachment 6-well plates (Corning-Costar) to induce spheroid formation. Spheroids were maintained for 6 days in NbM supplemented with 0.5 µM RA and 0.1 µM SAG. Spheroids were dissociated into single cells using dissociation solution (1:1 0.25% Trypsin-EDTA & Accumax) (ThermoFisher) and maturation of sMN was carried out under adherent conditions (as described above) in NbM supplemented with 0.5 µM RA, 0.1 µM SAG, 0.1 µM Compound-E (StemCell Technologies), 2ng/mL BDNF, 2ng/mL GDNF, and 2ng/mL CNTF (Life Technologies). After 72 h, NbM was replaced and additionally supplemented with 0.02 µM SN38-P to purify cultures of proliferative progenitors and undifferentiated stem cells (Mao et al., 2018). sMN were cultured under these conditions for a further 4-9 days.

### DHMN1 SV genotyping PCR assay

DNA was isolated from patient and control iPSC lines using the QuickExtract DNA Extraction Solution 1.0 (Epicenter Bio) according to the manufacturer’s instructions. Multiplex PCR amplification was performed in a 10 μL reaction containing 1X MyTaq HS Red Mix (Bioline), 10 ng DNA template, 4 pmol each of the forward and reverse primers and 8 pmol of the dual forward/reverse (multiplex) primer. Water was used as a blank control. Thermal cycling conditions and primers have been described previously ^2^. Agarose gels (1.5%) were prepared in 1 X TAE buffer (Astral Scientific) with 0.01% SYBR Safe DNA gel stain (Bioline). PCR products (5 µL) were size fractionated for 30-35 min at 90 V (40 V cm^-1^). Gels were visualised using a Safe Imager Transilluminator 2.0 (Invitrogen) and images captured using a Canon PhotoShot S5-IS digital camera with Hoya O(G) filter.

### *UBE3C-IF* PCR and sanger sequencing

Total RNA was isolated from patient and control sMN using the RNeasy mini kit (Qiagen) according to the manufacturer’s instructions. Yields were assessed using Nanodrop (Thermofisher). RNA (0.5 µg) was reverse transcribed using the iScript cDNA Synthesis kit (BioRad) in accordance with the manufacturer’s instructions. PCR amplification was performed in a 10 μL reaction containing 1X MyTaq HS Red Mix, 25 ng cDNA template and 4 pmol each of the forward and reverse primers. Water was used as a blank control. Thermal cycling was performed as previously described.^2^ PCR products were size fractionated by agarose gel electrophoresis and gels imaged as described above. For sequence validation, PCR products (5 µL) from samples P1 and C1, as well as 10 µM (10 µL) forward and reverse primers were provided to the Garvin Molecular Genetics Facility, Garvan Institute of Medical Research (Sydney, Australia) and sequencing was performed using BigDye Terminator Cycle Sequencing protocols. Electrophoresis primer pair: Forward – 5’-GGTACCCAAAGTCAGGAAGC-3’, Reverse – 5’-CAAAGCAGCAGTTCGAGTCT-3’. Sequencing primer pair: Forward – 5’ CTGCTCAACCTGGTGTGGA-3’, Reverse – 5’-CAAAGCAGCAGTTCGAGTCT-3’.

### *UBE3C-IF* qualitative RT-PCR (qRT-PCR)

Total RNA was isolated from DHMN1 and control tissues and reverse transcribed as described above. Quantitative RT-PCR was performed in 20 µL reaction volumes containing 1X TaqMan Gene Expression (Applied Biosystems) Assay, 1X TaqMan Gene Expression Mastermix (Applied Biosystems) and 50ng cDNA template. Water was used as a negative control in all assays. Thermal cycling was performed with a StepOnePlus Real-Time PCR machine (Applied Biosystems) using the following cycling protocol: 95°C for 20 s; followed by 40 cycles of 95°C for 1 s and 60°C for 20 s. *GAPDH* was used as an internal housekeeping gene. The control sMN group was used as the reference group. TaqMan Gene Expression probes used in this study can be found in supplementary (Supplementary Table S1).

### Immunohistochemistry

Cells were washed once in DPBS (Gibco), fixed in 4% (v/v) paraformaldehyde (PFA, Sigma) for 20 min at room temperature, washed once with DPBS, treated with permeablisation solution (DPBST) containing DPBS and 0.3% (v/v) Triton X-100 (Calbiochem) for 30 min at room temperature and blocked (DPBS- and 5% (w/v) bovine serum albumin (Sigma)) for 1 h at room temperature. Cells were incubated with primary antibodies overnight at 4°C. iPSC: anti-OCT4 (Cell Signalling, #2840, 1:400), anti-SOX2 (Cell Signalling, #3579, 1:400), and anti-NANOG (Cell Signalling, #4903, 1:400). MNP: anti-OLIG2 (Millipore, MABN50 1:100). sMN: anti-MNX1 (Sigma, HPA071717, 1:500), anti-68kDa NFL (Abcam, ab24520 1:1000) and anti-TUBB3 (Sigma, T2220, 1:1000). Details of all primary antibodies used in this study can be found in supplementary (Supplementary Table S2).

The cells were then incubated with Alexa Fluor (AF) secondary antibodies (Invitrogen) for 3 h at room temperature. The following secondary antibodies were used: AF goat anti-rabbit (488 or 555, 1:500), AF goat anti-mouse (488 or 555, 1:500) and AF goat anti-chicken (647, 1:500). Nuclei were counter-stained with 4,6-diamidino-2-phenylindole (DAPI, Molecular Probes, 1:5000) and mounted using Prolong Gold Antifade reagent (Invitrogen). Samples were imaged on a Leica SP8 confocal microscope and visualised using LAX software (Leica Microsystems). Images were processed using FIJI (version 2.0) for Mac OSX.

### Quantification of MNX1 stained nuclei

sMN positive for MNX1 were quantified using CellProfiler (version 3.1.9). DAPI-stained nuclei were segmented via “Otsu”-based global thresholding and used to define regions of interest (ROI) for the measurement of fluorescence of nuclei positive for MNX1. Global thresholding using the “Minimum cross entropy” method was applied to the MNX1 channel to determine a fluorescence value for positively stained nuclei. The “RelateObjects” function was used to overlay MNX1 images on their corresponding DAPI images. The percentage of positive nuclei was determined by dividing the number of MNX1^+^/DAPI^+^ co-positive nuclei by the number of DAPI^+^ nuclei. Data were obtained for each line from three independent differentiations.

### Quantification of neurite occupied area

Quantification was performed using CellProfiler (version 3.1.9). Images of TUBB3-stained axons were transformed to greyscale and segmented via “Otsu”-based global thresholding. Grey values (intensity) for each pixel were determined for each image using the “MeasureImageAreaOccupied” function. Area occupied by neurites was calculated by dividing grey pixels by the total number of pixels per image. Value is expressed as a percentage. Data were obtained for each line from three independent differentiations.

### Western blot

Samples were lysed using 1 X radioimmunoprecipitation assay (RIPA) buffer (Merck) supplemented with 1X cOmplete EDTA-free protease inhibitor (Roche). Protein concentrations for each sample were analysed using Pearson’s Bicinchoninic Acid (BCA) protein analysis kit (Thermofisher Scientific) in accordance with the manufacturer’s protocols and measured on an EnSpire Multimode Plate Reader (Perkin Elmer). Western blot analysis was carried out using 20 µg protein lysate. Protein samples were size fractionated on 4-15% Mini PROTEAN TGX Precast Gels (BioRad) via SDS-polyacrylamide gel electrophoresis at 130 V for 1.5 h in running buffer (25 mM Tris, 192 mM glycine, SDS-PAGE 0.1 % SDS, pH 8.3) and then transferred to Immobilin-P polyvinylidene difluoride (PVDF) transfer membranes (Merck) at 70 V for 75 min in transfer buffer (BioRad). Membranes were blocked (5% w/v skim milk powder (Oxoid) in 1 X Tris-Buffer Saline (TBS)) for 1 h at room temperature followed by overnight incubation at 4°C with primary antibodies prepared in TBS-T (TBS, 0.1% v/v Tween20). Primary antibodies: MNX1 (Sigma Aldrich, HPA071717, 1:500), UBE3C (Invitrogen, #PA5-110540, 1:500). Membranes stained with Ponceau S (Sigma Aldrich) were used to control for protein loading in these experiments. Details of all primary antibodies used in this study can be found in supplementary (Supplementary Table S2). The membranes were then incubated in anti-rabbit (Sigma Aldrich) and anti-mouse (Abcam) horseradish peroxidase (HRP) conjugated secondary antibodies (1: 5000) and signal detected using Clarity^TM^ Western ECL Substrate solution (BioRad). Blots were visualised on a ChermiDOC XRS+ system (BioRad) and images processed using Image Lab software v5 (BioRad). Intensity of protein bands were quantified using FIJI (version 2.0).

### NanoString nCounter gene expression assay

Gene expression quantification using the NanoString nCounter gene expression system (NanoString Technologies Inc) was outsourced to Westmead Medical Research Institute (Sydney, Australia). A 72-target custom panel (63 gene targets, 2 MNP markers, 2 sMN markers and 5 housekeeping genes) was designed to include all candidate genes 3 Mb on either side of the DHMN1 insertion breakpoints and genes within the DHMN1 complex insertion. The target-specific oligonucleotides were designed by NanoString Technologies (Washington, USA) and synthesized by Integrated DNA Technologies (Iowa, USA). Total RNA was isolated (sMN: D12, n = 5; MNP: D6, n = 4; iPSC: D4, n = 3) from patient and control tissues using the RNeasy mini kit (Qiagen) and diluted to a concentration of 10ng/µL. RNA yield was assessed using Nanodrop and integrity determined using TapeStation (Agilent). Gene count normalisation and expression analysis was performed using the nSolver Analysis Software package (version 4.0) (NanoString Technologies). The normalisation of the gene count data was performed using the recommended parameters described in the nSolver User Manual (Version 4.0). Background correction and low count gene filtering was performed via the in-built background thresholding function using the default threshold value of 20 counts. Details regarding assay preparation and execution can be found in Supplementary Methods.

### RNA sequencing data processing and analysis

sMN total RNA from DHMN1 patient (*n* = 3) and controls (*n* = 3) was isolated using the RNeasy mini kit (Qiagen) according to the manufacturer’s instructions. Yields were assessed using Nanodrop and RNA integrity determined using 2100 Bioanalyser (Agilent). Library preparation and mRNA sequencing was outsourced to Macrogen (Seoul, South Korea). In brief, polyA species selection and library preparation was carried out using the TruSeq Stranded mRNA LT Sample Prep Kit (Illumina). Paired-end (151 bp) sequencing was performed on an Illumina NovaSeq 6000 sequencer. Quality control (QC) was performed by Macrogen using FastQC^49^ with adapter sequences and low-quality bases trimmed using Trimmomatic.^50^ Trimmed reads were mapped to GRCh38/hg38 using the splice-aware aligner HISAT2.^51^ Quantification of gene and transcript level abundances and the identification of novel and alternative splicing transcripts was carried out using StringTie.^52^ Identification of fusion gene products was performed using Defuse^53^, FusionCatcher^54^ and Arriba^55^ programs. Gene-level differential gene expression analysis was performed using DESeq2.^56^ Differentially expressed genes (DEGs) were determined by log_2_FC ≥ abs 1.0 and a false discovery rate (FDR) adjusted *p-*value threshold of 0.05. Gene Ontology and enrichment analysis to determine biologically relevant pathways was performed using the Database for Annotation, Visualisation and Integrated Discovery (DAVID)^57,58^ and StringDB.^59–62^

### Chromatin Conformation Capture and paired-end sequencing (Hi-C)

Hi-C library preparation and sequencing was outsourced to Dovetail Genomics (Santa Cruz, California). A library was prepared from motor neurons from a DHMN1 patient (*n* = 1) and sequenced (150 bp paired-end) using three lanes of an Illumina HiSeq X to generate ∼300 Gb per lane. This would generate 900 million reads for a 10 kb binsize resolution (non-overlapping sequences) in which more than 80% of all possible bins would have 1000 or more reads (contacts). Mapping of Hi-C libraries and normalisation was performed using HiC-Pro pipeline.^63^ Sequencing reads were mapped to the hg38 reference genome and a hg38 custom-built DHMN1 chromosome 7 with the 1.35 Mb insertion, using the Bowtie aligner and assessed for artefact levels using the human genome and restriction enzyme cut sites. BAM files mapping the paired-end tags (PETs) were filtered to eliminate invalid PETs. Visualisation and analysis of genome-wide contact matrices from mapped and normalised Hi-C PET data was performed using HiGlass.^64^ Visualisation, identification, and prediction of TADs was performed using TADtool.^65^

### Construct generation and cloning

Q5 Site Directed Mutagenesis (SDM; New England Biolabs) was used to generate the pCMV6-Entry-UBE3C-IF plasmid. This was conducted in two stages. (1) the commercial template plasmid pCMV6-Entry-UBE3C (Origene #:RC215110) was amplified using primers which partially inserted the intergenic fusion sequence and simultaneously deleted the UBE3C coding sequence downstream of p.Val443 (F: 5’-aagaggatcattACGCGTACGCGGCCGCTC-3’; 5’-gaattgcttcctGACTTTGGGTACCATCATGCGGTGCT G-3’). (2) Q5 site-directed mutagenesis was again conducted on the plasmid generated in step 1 to introduce the remaining intergenic fusion sequence alone (5’-caccagaataaaACGCGTACGCGGCCGCTC-3’; 5’-acatcttgttaaAATGATCCTCTTGAATTGCTTCCTGACTTTGGGTACC-3’). The UBE3C-IF coding sequence was then amplified from pCMV6-Entry-UBE3C-IF using primers with flanking *Xba* I and *Not* I restriction sites (F: 5’-aaaaaatctagaATGTTCAGCTTCGAAGGC-3’; R: 5’-aaaaaagcggccgcCTATTTATTCTGGTGACATCTTGTTAAAATG-3’). This amplicon was then inserted into the pPD157.60 vector containing the unc-25 promoter sequences using the *Xba* I and *Not* I restriction sites to form the expression plasmid pPD157.60-UBE3C-IF. The ISOLATE II Plasmid mini (Bioline) was used for the purification of expression plasmids according to manufacturer’s instructions.

### Overexpression of *UBE3C-IF* in HeLa cells

HeLa cells (1 x 10^6^) were seeded onto 6-well plates and maintained in H-DMEM at 37°C and 5 % CO_2_ for 24 h. Cells were transfected with either empty vector or the expression plasmid pCMV6-Entry-UBE3C-IF (0.5 µg, 1.0 µg, 2.0 µg) via lipofection using Lipofectamine 3000 (Invitrogen) as per the manufactures protocol. Cultures were harvested 48 h post-transfection for western blot analysis as described above.

### C. elegans methods

Transgenic *C. elegans* strains were generated by microinjecting a cocktail of the expression plasmid containing UBE3C-IF or the empty vector at a concentration of 50 ng/µL and plasmid PCFJ90, a co-injection marker, at a final concentration of 5 ng/µL. Three independent transgenic lines were generated, and the experiments were carried out using the line that stably inherited the human transgene.

Strain information and additional *C. elegans* methods can be found in Supplementary Methods.

### Statistical analysis

For the statistical analysis of NanoString gene expression data, differential expression and significance was determined using the nSolver software (v4.0). The Differential Expression Call (DE Call) test function was used to predict differential expression for MNP and iPSC data where biological replication was limited. Statistical analysis of qRT-PCR data was carried out using the StepOne Plus Software (v2.1) (Applied Biosystems). For the immunofluorescence quantification data, significance was determined using a two-tailed Student’s t-test. Statistical analysis of RNA-seq was performed in RStudio utilising a custom DESeq2 pipeline. The significance of differential gene expression was determined using the Wald Test assuming a log_2_FC threshold of abs 1.0 and a false discovery rate (FDR) adjusted *p-*value threshold of 0.05. The statistical analysis and significance of western blot experiments was determined using either a two-tailed Student’s t-test (sMN experiments) or a one-way ANOVA followed by Dunnet’s multiple comparison test (HeLa experiments). The following statistical thresholds were applied throughout the study: \**p* < 0.05; \*\**p* < 0.01; \*\*\**p* < 0.001; \*\*\*\**p* < 0.0001.

### Data availability

All relevant data are included within this manuscript and supplementary material files. Raw RNA-seq data will be available upon reasonable request.

## Results

### Characterisation of DHMN1 patient-derived iPSC lines

Skin fibroblasts from three DHMN1 patients were reprogrammed by FUJIFILM Cellular Dynamics International (CDI) using non-integrative episomal plasmids and company in-house protocols. To fully validate the iPSC lines, in-house pluripotency analysis was performed, and molecular analysis confirmed the DHMN1 complex insertion was retained after the reprogramming process. These results were confirmed in the iPSC lines from three different patients and representative data is presented (Fig. 1). Normal karyotyping was observed (Fig. 1A) and genotyping DNA extracted from the patient-derived iPSC lines confirmed the presence of the SV mutation post-reprogramming (Fig. 1B). Pluripotency of the DHMN1 iPSC lines was initially confirmed by CDI as part of the service. Subsequent immunofluorescence experiments in our laboratory also confirmed the DHMN1 iPSC lines were positive for the pluripotency markers OCT4A, SOX2 and NANOG (Fig. 1C).

**Figure 1:**
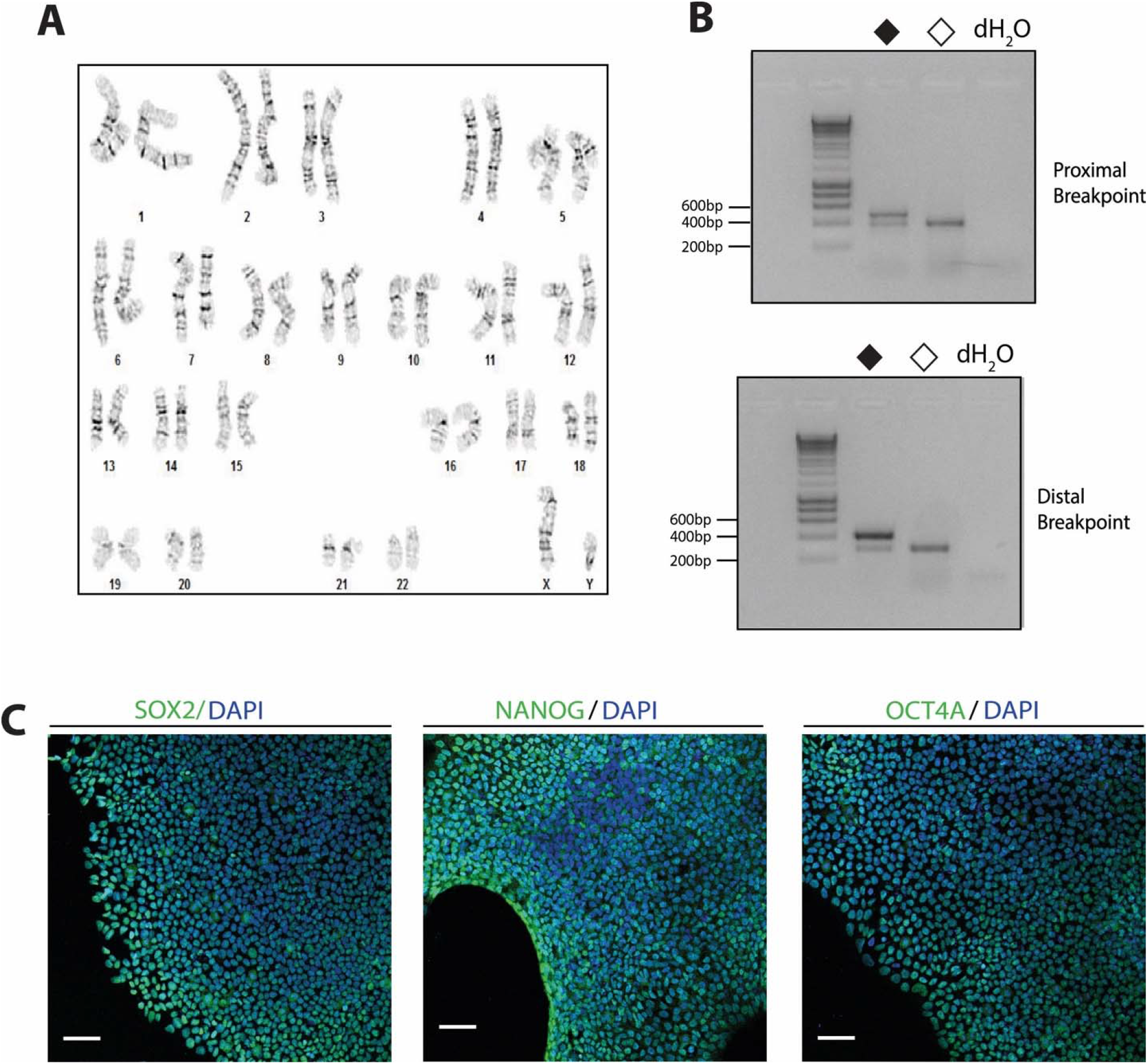
DHMN1 iPSCs harbouring the 1.35 Mb complex insertion display a normal karyotype and exhibit molecular features of pluripotency. (**A**) Karyotyping and G-band analysis confirm that iPSC lines have a normal 46, XY karyotype. (**B**) DHMN1 multiplex breakpoint genotyping assay for a representative patient (P1, closed diamond) and control (C1, open diamond) confirms the presence of the complex insertion in DHMN1 iPSC lines. The mutant junction fragments (501 bp, proximal; 412 bp, distal) and wild-type fragments (409 bp, proximal and 290 bp distal) were amplified from the respective DHMN1 and wild type chromosomes. (**C**) Immunofluorescence staining confirms expression of pluripotency markers OCT4, SOX2 and NANOG for a representative patient iPSC sample (P2). Images are presented as pluripotency markers (green) overlayed on DAPI stained nuclei (blue). Scale bar 60 µm.

### Generation of MNP and sMN by differentiation of DHMN1 patient-derived iPSC lines

To develop a motor neuron model for DHMN1, 3 patient and 3 control iPSC lines were differentiated into sMN from highly expandable populations of *OLIG2^+^* positive motor neuron progenitors (MNP) as initially reported by Du *et al*.^48^ Generating sMN using MNP provided added practicality as they can be cryopreserved, and undergo thawing for differentiation, thereby reducing the overall time to produce sMN. In addition, culturing MNP produces highly pure populations of sMN, and avoids many of the pitfalls associated with other differentiation methods including poor yields and low efficiency^48^. We modified the protocol to include a purification step whereby adherent single cell sMN were cultured in the presence of 0.02 µM SN38-P for 96 h (Fig. 2A). The SN38-P acts to purify the culture of any remaining proliferative, undifferentiated iPSC and MNP^66^. We have previously shown SN38-P treatment enriches cultures for mature sMN without the need for additional sorting protocols^67^. Immunohistochemistry confirmed that iPSC differentiated into ventralised neural stem cells expressing OLIG2 (MNP, Supplementary Fig. S1A). Additionally, the NanoString data demonstrated that *Nestin* (*NES),* also a marker of neural stem cells^68^ was expressed at higher levels in differentiated cells compared to iPSC (Supplementary Fig. S1B). Together these data indicate successful differentiation of iPSC into MNP.

**Figure 2:**
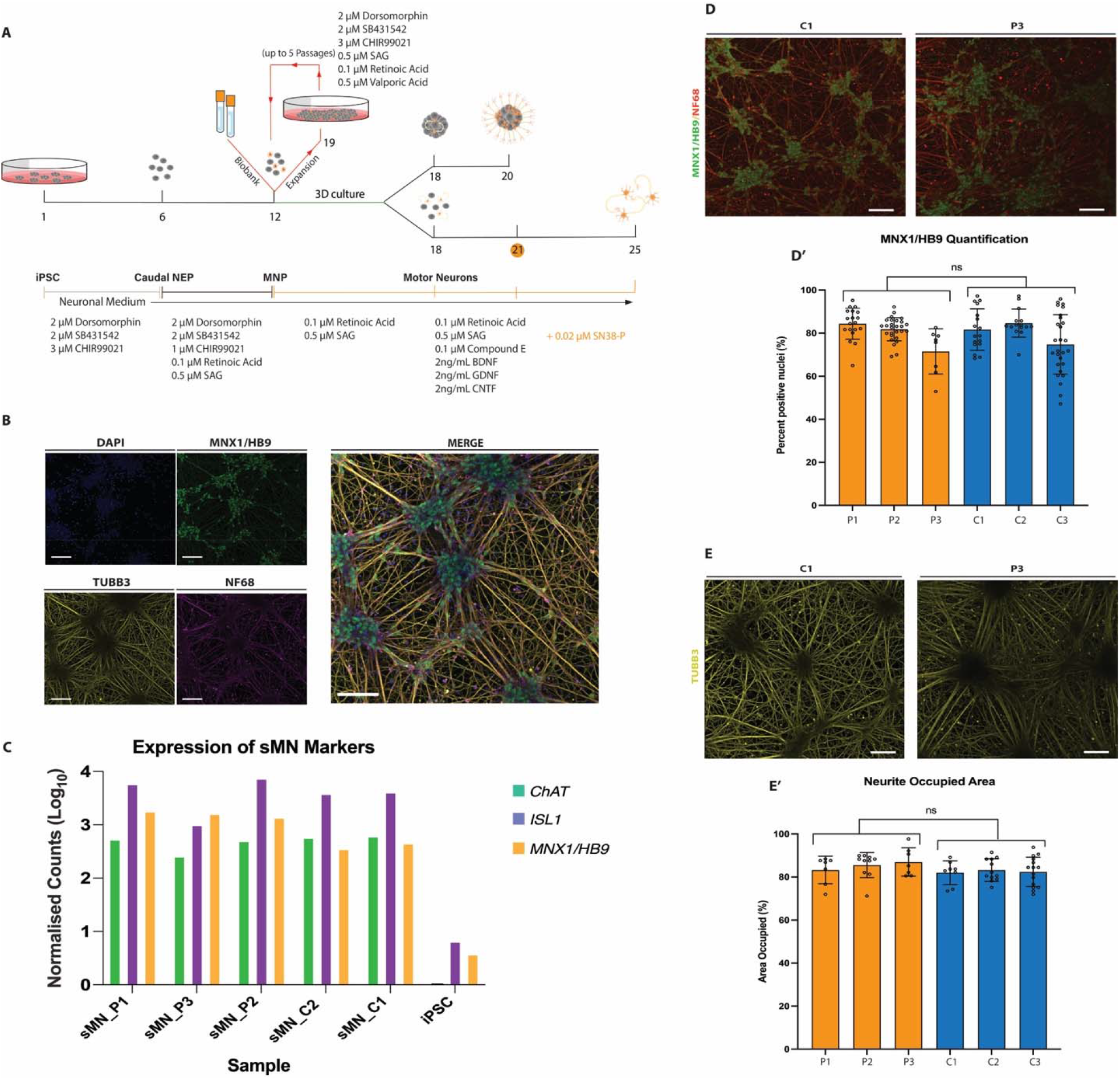
sMN derived from patient and control iPSC do not show changes in number of MNX1/HB9^+^ sMN or 2D neuronal networks. **(A)** Modified differentiation workflow used to generate MNP and sMN. Timeline of the differentiation process, media and factors used throughout the protocol. The yellow circle at day 21 indicates initiation of treatment with SN38-P **(B)** Immunofluorescence staining of iPSC-derived sMN for MN markers TUBB3, MNX1/HB9 and NF68. Scale bar 90 µm. **(C)** NanoString measurements comparing sMN and iPSC show increased expression of markers for motor neuron identity *ISL1, MNX1/HB9* and *ChAT*. **(D)** Immunofluorescence staining of a representative patient and control sMN line positive for MNX1/HB9 and NF68. Scale bar 90 µm. **(D’)** Comparison of the number of sMN generated from patient (*n* = 3) and control (*n* = 3) lines. Approximately 2500 cells were analysed per line. Data was pooled for each line from three independent rounds of differentiation. The number of sMN are represented by the percentage of nuclei stained for MNX1/HB9. **(E)** Immunofluorescence staining of a representative patient and control sMN line positive for TUBB3. Scale bar 90 µm. **(E’)** Quantification of TUBB3 fluorescence in patient (*n* = 3) and control (*n* = 3) sMN lines reveals no difference in 2D neuronal networks. Data was pooled for each line from three independent rounds of differentiation.

MNP that were cultured in suspension for 6 days formed 3D aggregates (neurospheres) of MNP actively differentiating into sMN. Following the disaggregation of these neurospheres, the sMN were matured under adherent 2D culturing conditions for an additional 7-12 days. The mature sMN demonstrated robust expression of the sMN marker MNX1, the neuronal cytoskeletal marker βIII-Tubulin (TUBB3) and the axonal marker NF68 (Fig. 2B). Similarly, the expression of the sMN markers *MNX1, ISL1* and *ChAT*, determined by the NanoString gene read counts, were higher in sMN when compared to the iPSC (Fig. 2C). Quantification of the MNX1^+^ nuclei revealed comparable numbers of sMN generated for the different patients and controls (Fig. 2D-D’). Similarly, quantification of TUBB3 did not reveal any differences in the 2D neuronal networks between the patients and controls (Fig. 2E-E’).

### Genes within and flanking DHMN1 insertion breakpoints are dysregulated

Initial linkage studies mapped the DHMN1 locus to a 12.3 Mb region at chromosome 7q34-q36.3^69^ containing 112 candidate genes. Identification of the SV mutation enabled prioritisation of possible candidate genes by 1) copy number, 2) disruption of the normal regulatory environment or, 3) introduction of new regulatory elements driving ectopic expression of nearby genes (see Kleinjan^70–72^ for review). To analyse the local gene expression signatures associated with the DHMN1 complex insertion, 63 candidate transcripts were selected for NanoString analysis in iPSC, MNP and sMN tissues (Supplementary Table S3). We hypothesised the genomic rearrangement causing the DHMN1 phenotype could affect genes lying within the DHMN1 complex insertion as well as genes flanking the insertion breakpoints – given that cis-regulatory control of gene expression has been reported to occur over distances of up to 1.45 Mb.^73^ The custom assay was therefore designed to select genes localising within the DHMN1 complex insertion or in the genomic regions (3 Mb) flanking the DHMN1 insertion breakpoints (Fig. 3A).

**Figure 3:**
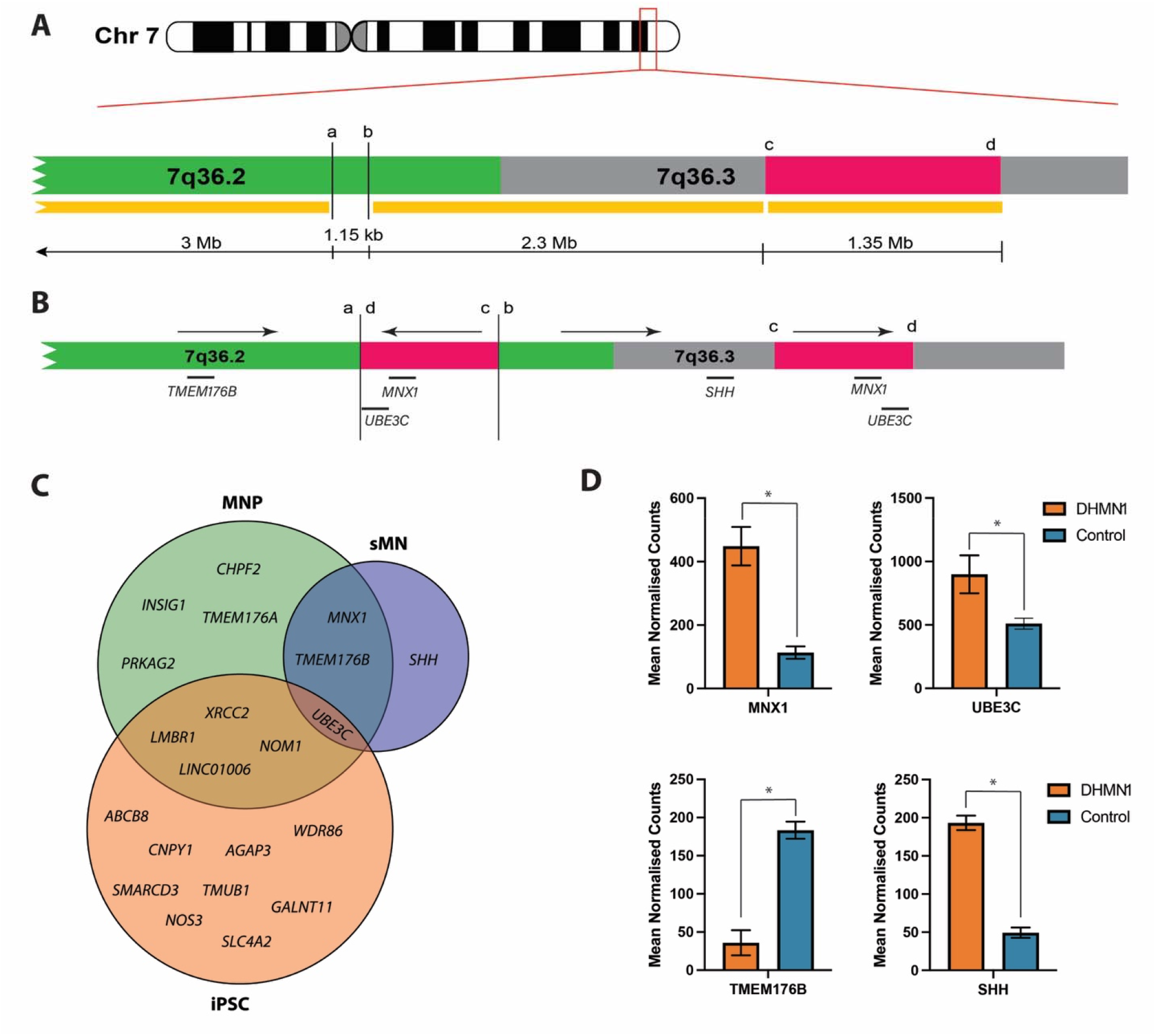
NanoString assay reveals nine differentially expressed genes (DEG) within a 6 Mb region flanking the DHMN1 insertion breakpoint sites. (**A**) Ideogram and localising the genomic region of chromosome 7q36.2-q36.3 involving the formation of the DHMN1 complex SV rearrangement. The green region represents the DHMN1 linkage region. The pink region represents the genomic interval at 7q36.3 defining the DHMN1 insertion sequence. Breakpoints are indicated by the vertical solid black lines. Letters a-d indicate genomic co-ordinates at the proximal (a) and distal (b) ends of the insertion site and the proximal (c) and distal (d) ends of the 7q36.3 duplicated sequence. Yellow bars indicate the genomic regions used to select genes for differential gene expression analysis with a custom designed NanoString panel. Horizontal black lines show genomic distances. **(B)** Schematic representation of the DHMN1 locus showing the relative positions of differentially expressed genes within the queried genomic intervals. Arrows indicate orientation of genomic sequences. Letters a-d indicate genomic co-ordinates at the proximal (a) and distal (b) ends of the insertion site and the proximal (c) and distal (d) ends of the 7q36.3 duplicated sequence which is now in an inverted orientation within the DHMN1 locus. The insertion breakpoints are denoted by the vertical solid black lines. **(C)** Venn diagram depicting the overlap of statistically significant DEG from iPSC, MNP and differentiated sMN. **(D)** Mean normalized counts for the four genes showing statistically significant DEG between DHMN1 patient (n=3) and control (n=2) sMN. Data is represented as mean ± standard error (SE).

Analysis of the expression signatures of the 63 genes showed that 10 targets and 14 targets were differentially expressed in MNP and iPSC respectively following background correction and filtering of low-count genes. (DE Call test; Supplementary Table S4). For the sMN, 4 of the 63 gene targets tested showed differential expression (DE) (*p* < 0.05) following background correction and filtering of low-count genes (Fig. 3D, Supplementary Table S5). For the four targets showing differential expression, two were within the DHMN1 insertion sequence (*MNX1, UBE3C*), one was within the DHMN1 linkage locus (*TMEM176B*) and one target was outside the DHMN1 locus and proximal to the insertion sequence originating from chromosome 7q36.3 (*SHH*) (Fig. 3B, Supplementary Table S3). The assessment of target genes in the MNP and iPSC displayed minimal overlap with the expression changes in sMN with *UBE3C* being the only differentially expressed gene across the three tissue types (Fig. 3C).

### The DHMN1 complex insertion alters 3D genome organisation

The structure and architecture of the 3D genome plays a crucial role in the spatiotemporal regulation of gene expression in tissues and cells. To assess how the 1.35 Mb complex insertion could change the genomic architecture and thereby impact downstream gene regulation at the DHMN1 locus, Hi-C analysis on patient (*n* = 1) and control (*n* = 1) sMN tissue was performed. Data was analysed using a custom, in-house, targeted Hi-C analysis pipeline. Topological associated domains (TAD) calling algorithms identified an altered TAD profile in patient sMN at the DHMN1 locus when compared to the control sMN profile (Fig. 4A). Visualisation and assessment of genome contacts across the DHMN1 locus also predicted aberrant genomic interactions occurring within patient sMN as well as the formation of a neo-TAD (Fig. 4B). Overlaying the contact matrices with the genomic map of the DHMN1 locus showed predicted aberrant interactions with the neo-TAD overlapping with the significantly dysregulated expression of the candidate genes *MNX1* and *UBE3C* (Fig. 4C). The observed congruence between the gene dysregulation and Hi-C data further prioritised these genes as potential causative DHMN1 candidates.

**Figure 4:**
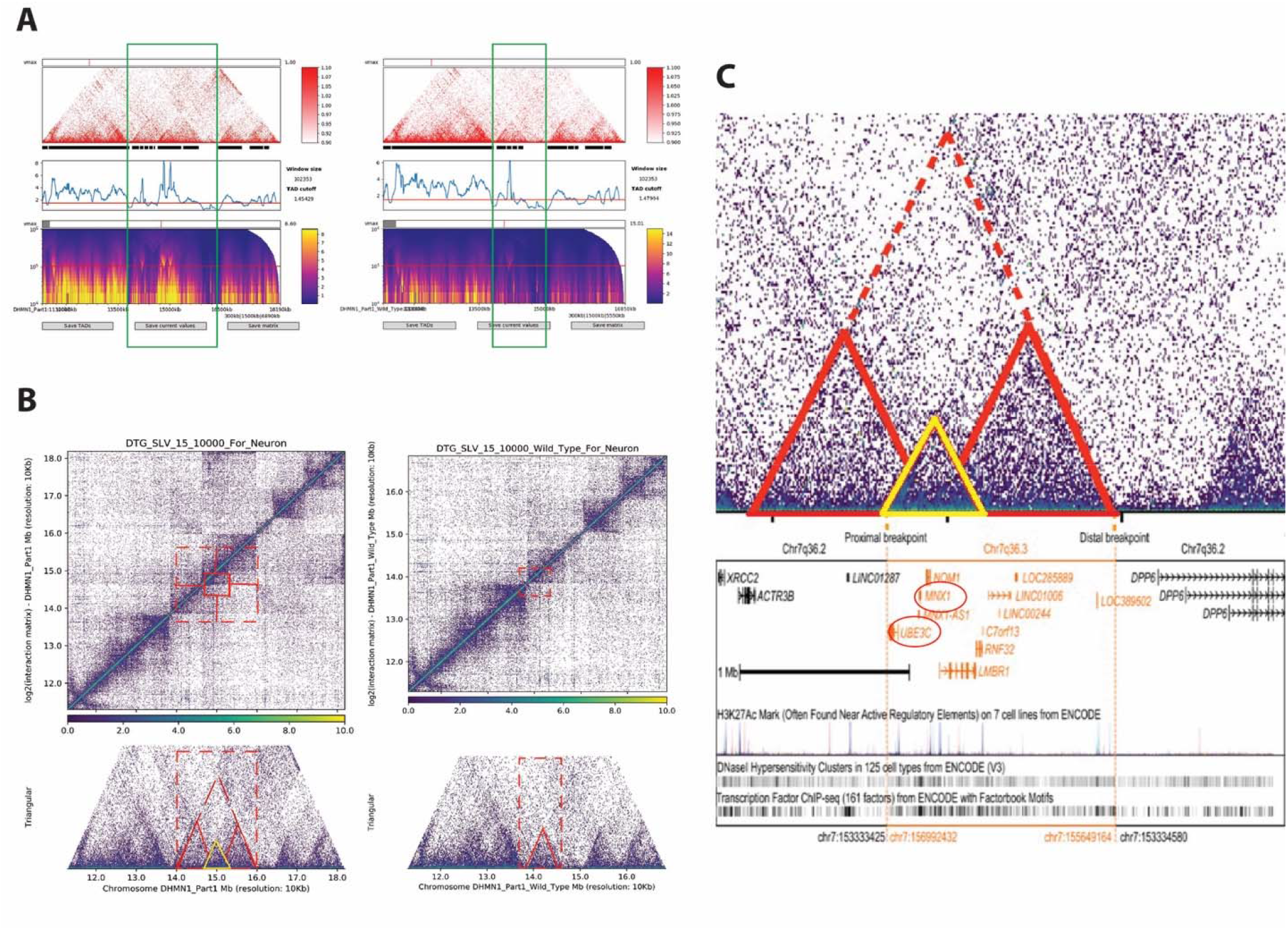
Hi-C analysis of DHMN1 sMN tissue identifies aberrant 3D genomic interactions and altered TAD profile at the DHMN1 locus in DHMN1 motor neurons. **(A)** Differential TAD profiles can be observed between patient (left panel, green box) and unaffected control (right panel, green box) sMN. **(B)** Contact matrices of DHMN1 (left panel, top and bottom) hg38 custom-built DHMN1 chromosome 7 and the hg38 reference genome (wild-type; right panel, top and bottom) sMN show 3D interaction profiles across the DHMN1 locus. Red boxes indicate differing regions of interaction. Red triangles depict the location of predicted TADs. A neo-TAD (yellow triangle) can be seen in patient sMN (left panel, bottom) suggesting the 1.35 Mb insertion changes the overall chromatin architecture at the DHMN1 locus creating new chromatin domains. **(C)** The Hi-C contact matrix when overlayed on a genomic map demonstrates the predicted neo-TAD (yellow triangle) overlaps with two significantly dysregulated candidate genes *MNX1* and *UBE3C* (red circles).

### RNA-seq excludes global gene dysregulation as a potential pathogenic mechanism and does not reveal biologically relevant pathways related to prioritised candidate genes

RNA-seq is an invaluable research tool enabling comprehensive quantitative analysis of the global transcriptome.^74,75^ One of the aims of this study was to compare global transcriptome profiles of DHMN1 and control sMN to elucidate genes, pathways and mechanisms that may be impacted by dysregulation of *UBE3C* or *MNX1*. To address this aim, RNA-seq was performed on polyA-selected mRNA transcripts extracted from patient (*n* = 3) and healthy control (*n*= 3) sMN cultures (day 12). Quality control, trimming and mapping statistics for each sample are summarised in supplementary data (Supplementary Table S6). An in-depth summary of data diagnostics is available in supplementary data (Supplementary Fig. S2 and S3). Following low count filtering, a total of 19,929 features were identified as expressed in all samples. Differential gene expression analysis of these expressed features revealed only 22 differentially expressed genes (DEGs) in DHMN1 sMN compared to healthy controls (Supplementary Table S7). The 22 genes did not localise to the DHMN1 locus and only one gene (*UNCX*) mapped to chromosome 7. The expression signatures of *UBE3C* and *MNX1* also showed the upregulation (Fig. S4 A and B) however, the changes did not reach statistical significance as observed in NanoString (Supplementary Fig. S4 A’ and B’). Gene Ontology (GO) analysis and functional annotation clustering was performed using StringDB and DAVID algorithms to determine the biological relevance of the 22 DEGs (Fig. 5A-D). After combining the output from both pipelines, a total of 30 significantly enriched terms were identified. The highest proportion of terms (13 out of 30) corresponded to tissue/organ development and morphogenesis (Fig. 5E). Of these, 50% (6 out of 12) involved the nervous system (Fig. 5F). No GO terms relating to neuronal compartments (axon, soma, dendrites), motor neurons, known pathways involved in CMT/IPN, motor neuron degeneration or other potentially biologically relevant pathways were identified across either analysis.

**Figure 5:**
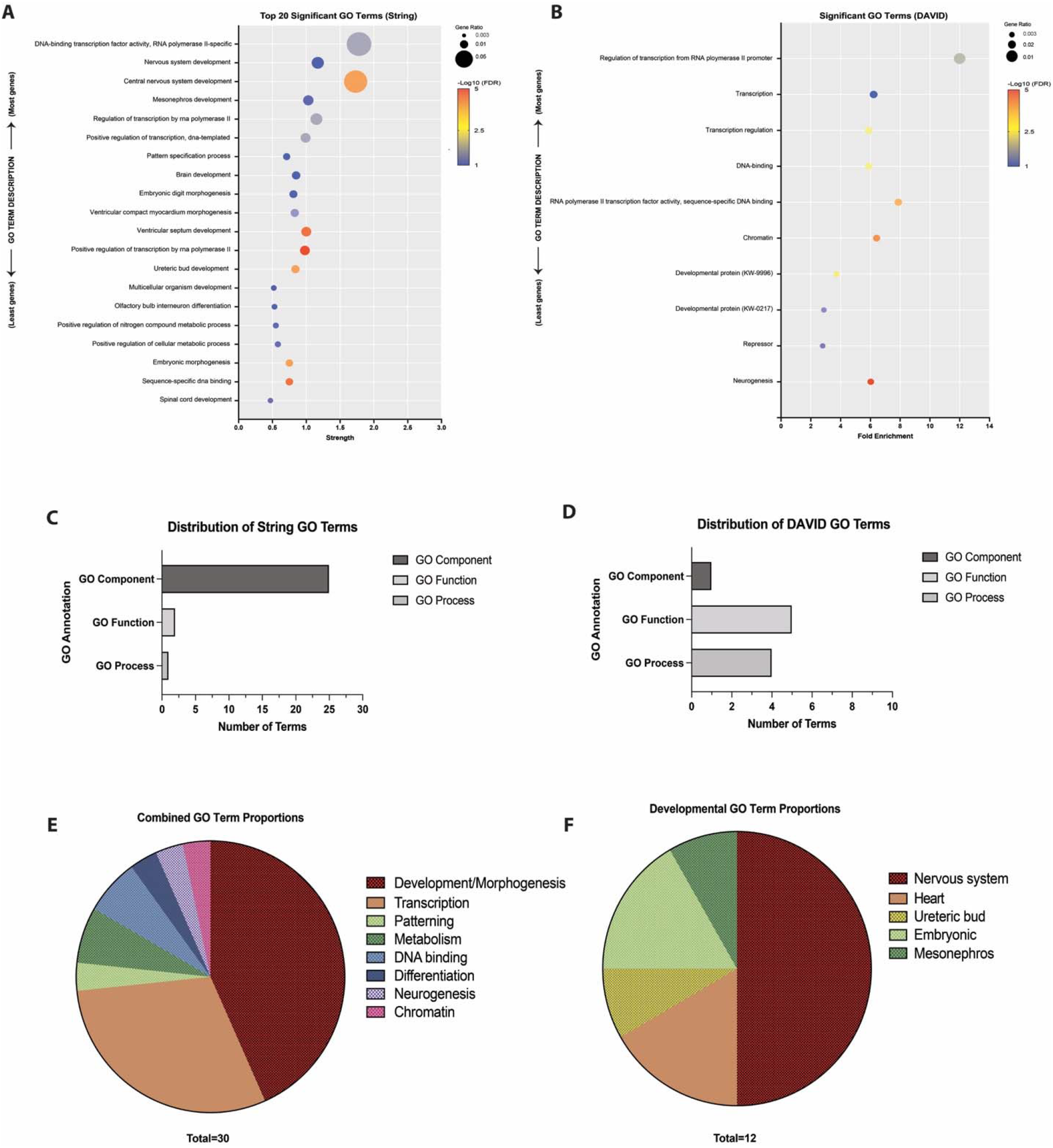
Gene ontology and enrichment analysis of the 22 DEGs identified in DHMN1 sMN shows no enrichment of biologically relevant terms, processes or pathways. Multiple variables plot for significant GO terms identified by **(A)** StringDB and **(B)** DAVID analysis methods. Terms are ranked according to those containing the most genes from the DEG list and plotted against the enrichment metrics for each method. The colour of each point represents the magnitude -Log10 P values and the size of each point represents the gene ratio (number of DEGs associated with a term vs the number of genes in the genome associated with the same term). **(C-D)** The breakdown of term categories for String DB and DAVID analysis methods. **(E)** The proportion of term annotations from the combined list of significant GO terms from DAVID and StringDB. **(F)** The breakdown of terms with annotations relating to development/morphogenesis.

### RNA-seq reveals a novel gene-intergenic fusion involving the *UBE3C* partial gene duplication

The DHMN1 SV involves the duplication and insertion of a 1.35 Mb region of chromosome 7q36.3 into the disease locus (chromosome 7q36.2) in an inverted orientation resulting in partial duplication (first 10 exons) of the *UBE3C* gene. Given that we have observed upregulation of *UBE3C* in DHMN1 sMN, we hypothesised that this was due to the production of a novel fusion transcript involving the partial *UBE3C*. The RNA-seq dataset was interrogated for segregating, predicted, novel gene fusions involving the partial *UBE3C* with a nearby gene or non-coding genomic sequence on the negative strand. A summary of chromosome 7 gene fusions can be found in supplementary material. The Arriba and Defuse algorithms detected a novel fusion involving the partial *UBE3C* sequence that was present in the three DHMN1 patient sMN samples (Fig. 6A; Supplementary Table S8). *In silico* splice site prediction analysis (NNSPLICE 0.99^76^) of the intergenic sequence downstream of the *UBE3C* partial gene strongly (score of 1.0 out of 1.0) indicated a canonical acceptor site ∼ 7 kb downstream of the final splice donor site of the partially duplicated *UBE3C* gene. The genomic sequence downstream of this splice acceptor site corresponded with the sequence of the novel terminal pseudo-exon (tPE) detected by Arriba and localised to intergenic DNA from the DHMN1 locus (Fig. 6B). RT-PCR and Sanger sequencing subsequently confirmed the presence of the *novel gene-intergenic fusion* sense transcript (*UBE3C-IF*) (Fig. 6B-C). To determine if a truncated *UBE3C-IF* was transcribed in addition to the wild-type full-length *UBE3C* (*UBE3C-WT*), quantitative RT-PCR using TaqMan probes was performed on sMN tissue from patients and controls. Significant differences in *UBE3C* expression were observed between patients and controls for a probe localising within the duplicated exons (*p* < 0.01). In contrast, no changes in expression between patients and controls were detected with a probe localising to the non-duplicated exons (Fig. 6D).

**Figure 6:**
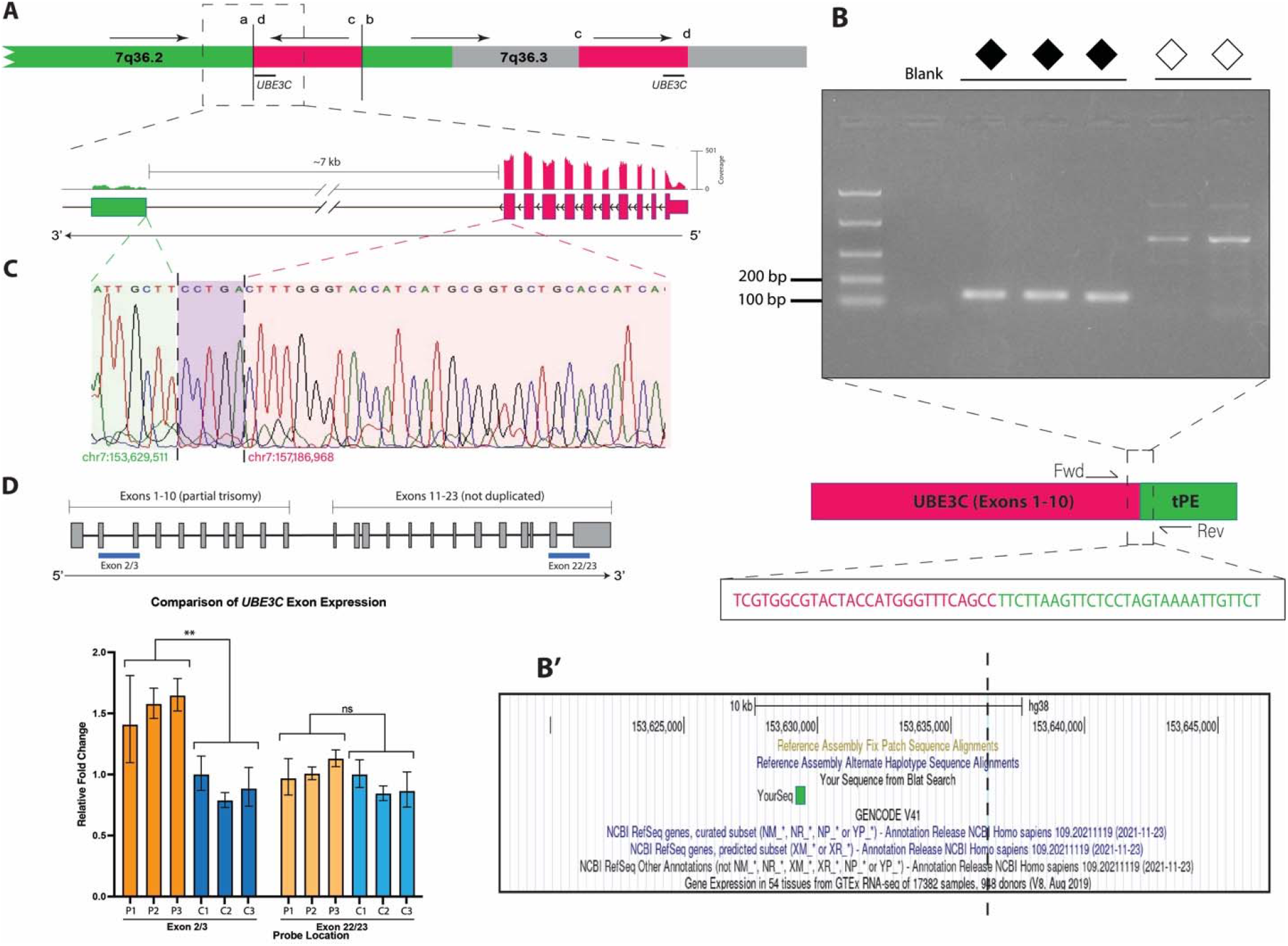
RNA-seq identifies a novel gene-intergenic fusion involving the partially duplicated *UBE3C*. **(A)** DHMN1 genomic rearrangement showing the depth of coverage of exonic sequences for the novel fusion transcript **(B)** RT-PCR validation of the *UBE3C* intergenic fusion transcript (*UBE3C-IF*) showing a 151 bp product amplified from the exon 10/pseudo-exon junction is present in patients (closed diamonds) and absent from controls (open diamonds). Expanded below to show a schematic of the fusion transcript. A terminal pseudo-exon (tPE) derived from intergenic DNA from within the DHMN1 locus is fused to exon 10 of *UBE3C*. **(B’)** BLAST of tPE sequence from RNA-seq (green box) aligns with 100% sequence identity to the intergenic region at chr7:153,629,188-153,629,514 in the DHMN1 locus. Dashed black line indicates insertion site. **(C)** Sanger sequencing of RT-PCR amplicons spanning the exon 10/pseudo-exon breakpoint confirms the presence of *UBE3C-IF* transcript in patient sMN mRNA. Coloured boxes indicate sequence alignments originating from the DHMN1 intergenic region (light green) and the partial *UBE3C* transcript exon 10 (light pink). The exact location of the end sequence from the partial *UBE3C* transcript exon 10 and start of the DHMN1 intergenic sequence cannot be unambiguously defined due to a 5 bp (CCTGA) overlap (purple box). **(D)** Schematic of the *UBE3C* transcript (refseq ID: NM_014671.1) showing duplicated and non-duplicated segments. Differential expression is observed between patient and control sMN when quantitative RT-PCR is performed using a TaqMan assay probe localising within the duplicated segment. Blue bars indicate probe location.

### DHMN1 sMN protein lysates show a reduction in full-length UBE3C

BLAST Protein^77–80^ confirmed the peptide sequence spanning the exon 10/pseudo-exon junction predicted by Arriba aligned to wild-type, canonical UBE3C-WT (NP_055486.2) with 100% sequence identity up to the pseudo-exon junction in all patients (Supplementary Fig. S5A-C). The full peptide sequence of UBE3C-IF is predicted to be 460 amino acids (aa) corresponding to a molecular weight of approximately 50.6 kDa (based on an average molecular weight of 110 Da/aa; Supplementary Fig. S5D). To investigate if *UBE3C-IF* is translated into protein, western blot analysis was performed on sMN protein lysates blotted with a UBE3C-WT antibody raised against the N-terminal of the protein (aa 1-270). Chemiluminescent visualisation revealed 2 additional bands below UBE3C-WT (123 kDa) at ∼ 55 kDa and ∼ 50 kDa (Fig. 7A). These bands may correspond to shorter protein isoforms of UBE3C that are annotated in Ensembl (Supplementary Fig. S6) and overlap with the predicted molecular weight of UBE3C-IF. Densitometric quantification of the two bands was performed to determine if the UBE3C-IF co-migrated with the shorter UBE3C isoform however, no difference was observed between the DHMN1 and control sMN tissues (Supplementary Fig. S7B-B’). Although it was not possible to resolve a specific band corresponding to UBE3C-IF, densitometric quantification of the full length UBE3C-WT protein showed levels that were significantly reduced in DHMN1 sMN tissue compared to controls (*p* < 0.0001; Fig. 7B). Western blot analysis of MNX1 (which was also prioritised as a candidate with differential gene expression) showed no difference between DHMN1 and control sMN tissue (Supplementary Fig. S7A-A’) and was therefore excluded as a pathogenic candidate gene.

**Figure 7:**
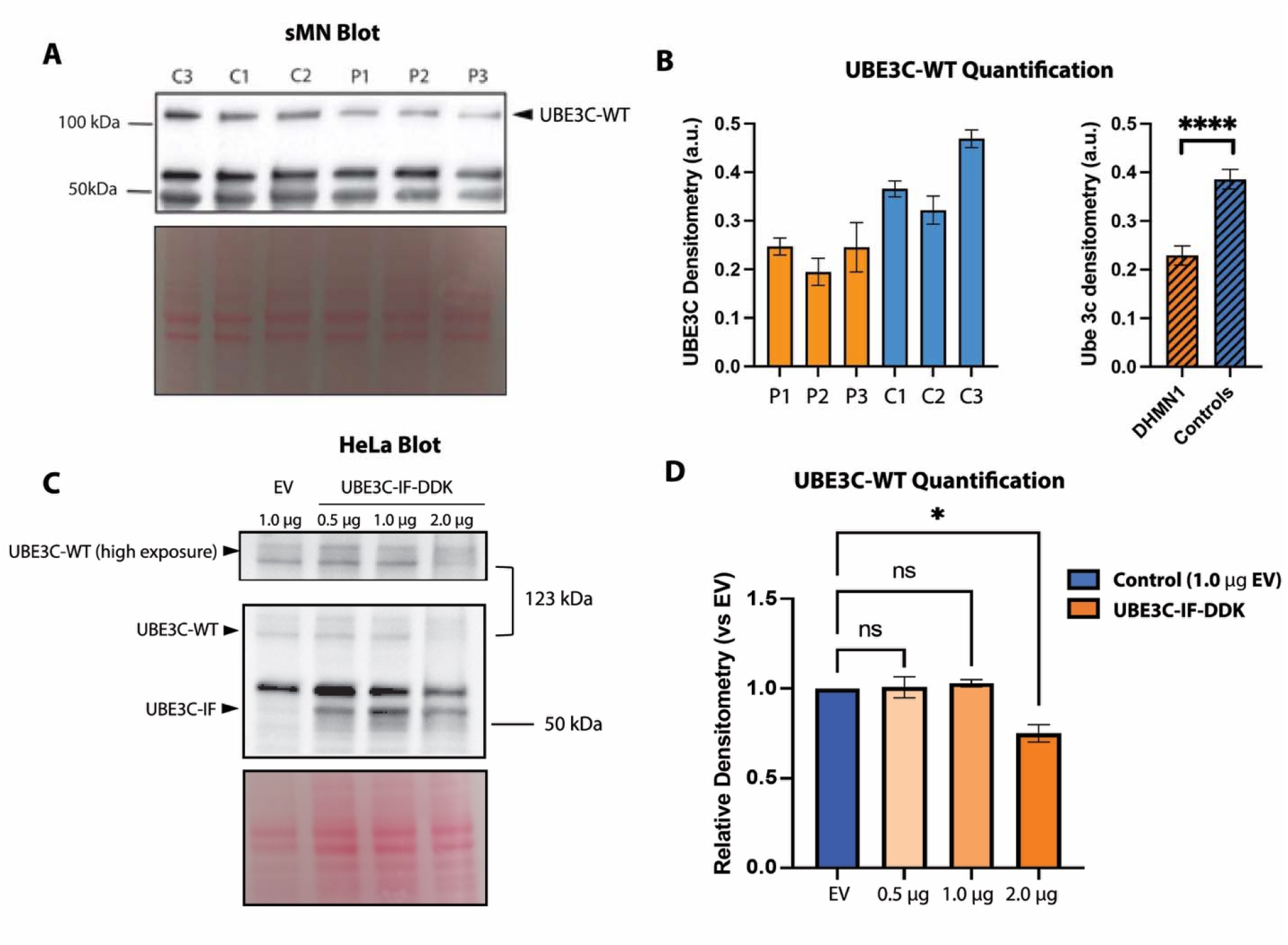
UBE3C-WT reduced in the presence of UBE3C-IF. **(A)** Representative western blot of UBE3C from sMN lysates. Ponceau stain is shown below (bottom panel). **(B)** Quantification of UBE3C-WT (123 kDa, top band) protein levels shows statistically significant reduction in DHMN1 sMN lysates compared to controls (*p* < 0.0001). UBE3C-WT was normalised to total protein (Ponceau). Quantification was performed using three independent experiments **(C)** A representative western blot of UBE3C-WT in HeLa cell lysates transfected with UBE3C-IF construct or empty vector (EV). UBE3C-IF can be detected using an antibody against UBE3C-WT (top band, middle panel). **(D)** Quantification of UBE3C-WT protein levels show a statistically significant reduction in cells transfected with 2.0 µg of UBE3C-IF construct when compared to EV (*p* < 0.05). Protein levels were normalised to Ponceau. Data is expressed as a ratio of UBE3C-WT expression in treatment conditions vs EV control. Quantification was performed using data from two independent experiments.

### Overexpression of *UBE3C-IF* in HeLa cells recapitulates reduction of full-length UBE3C-WT observed in patient sMN

To determine if the presence of UBE3C-IF results in the reduction of UBE3C-WT protein levels, HeLa cells were transfected with 0.5 -2 µg of construct containing *UBE3C-IF* (or empty vector; EV) and endogenous UBE3C was analysed from protein lysates harvested for western blot analysis using the UBE3C-WT antibody. Chemiluminescent visualisation of the blot showed cells transfected with the UBE3C-IF construct displayed an additional band corresponding to ∼ 50 kDa. This demonstrated that UBE3C-IF can be detected with an antibody raised against the N-terminal of the UBE3C-WT. Furthermore, this indicates that UBE3C-IF could co-migrate with a shorter, tissue-specific 50 kDa isoform of UBE3C observed in the sMN lysates (Fig. 7C). Whilst there was not a trend of decrease in endogenous UBE3C-WT protein levels with increasing amounts of UBE3C-IF construct, densitometric quantification showed that the cells transfected with 2.0 µg of UBE3C-IF had significantly reduced levels of UBE3C-WT (Fig. 7D) when compared to cells transfected with empty vector suggesting perturbed autoregulation at higher concentrations of UBE3C-IF.

### Overexpression of *UBE3C-IF* in *C. elegans* affects synaptic transmission, causes susceptibility to heat stress but does not affect neuronal morphology or locomotion behaviour

Aldicarb (an acetylcholine esterase inhibitor) and levamisole (an acetylcholine receptor antagonist) are drugs commonly used for screening *C. elegans* mutants defective in synaptic transmission.^81^ Using this method, we have previously successfully identified synaptic transmission deficits in a CMT model of *C. elegans.*^82^ *UBE3C-IF* animals displayed resistance to aldicarb induced paralysis when compared to control animals (Fig. 8A). In the presence of 1mM aldicarb, the time taken for 50% of animals to paralyse was approximately 70 min for *oxIs12* and *oxIs12;EmptyVector* when compared to 90 min for *oxIs12;UBE3C-IF* animals. Transgenic animals were hypersensitive to levamisole when compared to *oxIs12* animals (Fig. 8B). However, there was no significant difference between animals carrying *UBE3C-IF* plasmid and animals carrying the empty vector backbone. This suggested the hypersensitivity to levamisole may be due to the plasmid copy number and not the *UBE3C-IF* transcript. Our results show that the post-synaptic component (muscle) is not involved in the aldicarb resistant phenotype observed in *UBE3C-IF* animals thus implicating pre-synaptic compartment (axonal) deficits. This reflects the axonal presentation observed in DHMN1 patients. *oxIs12* animals carrying the transgene *unc-47::GFP,* allow visualisation of motor neurons in live *C. elegans*. Live imaging of age-synchronised day1 old *UBE3C-IF* and control animals showed intact axon and cell bodies with no signs of neurodegeneration (Fig. 8C and D). Correspondingly, the thrashing assay used to identify locomotion deficits associated with *UBEC-IF* overexpression showed no significant changes in locomotion between transgenic and control animals (Fig. 8E). Loss of *Hul5*, the yeast ortholog of *UBE3C*, results in reduced recovery rate following heat shock.^83^ To determine if the truncated *UBE3C-IF* construct could negatively impact a heat-induced protein unfolding response (the heat shock assay), *UBE3C-IF* and control animals were exposed to heat stress (8 h at 35°C). *UBE3C-IF* animals showed reduced survival rate (∼50%) when compared to controls animals [*oxIs12* (∼82%) and *oxIs12;EmptyVector* (∼77%)] (Fig. 8F) suggesting reduced fitness in response to protein unfolding.

**Figure 8:**
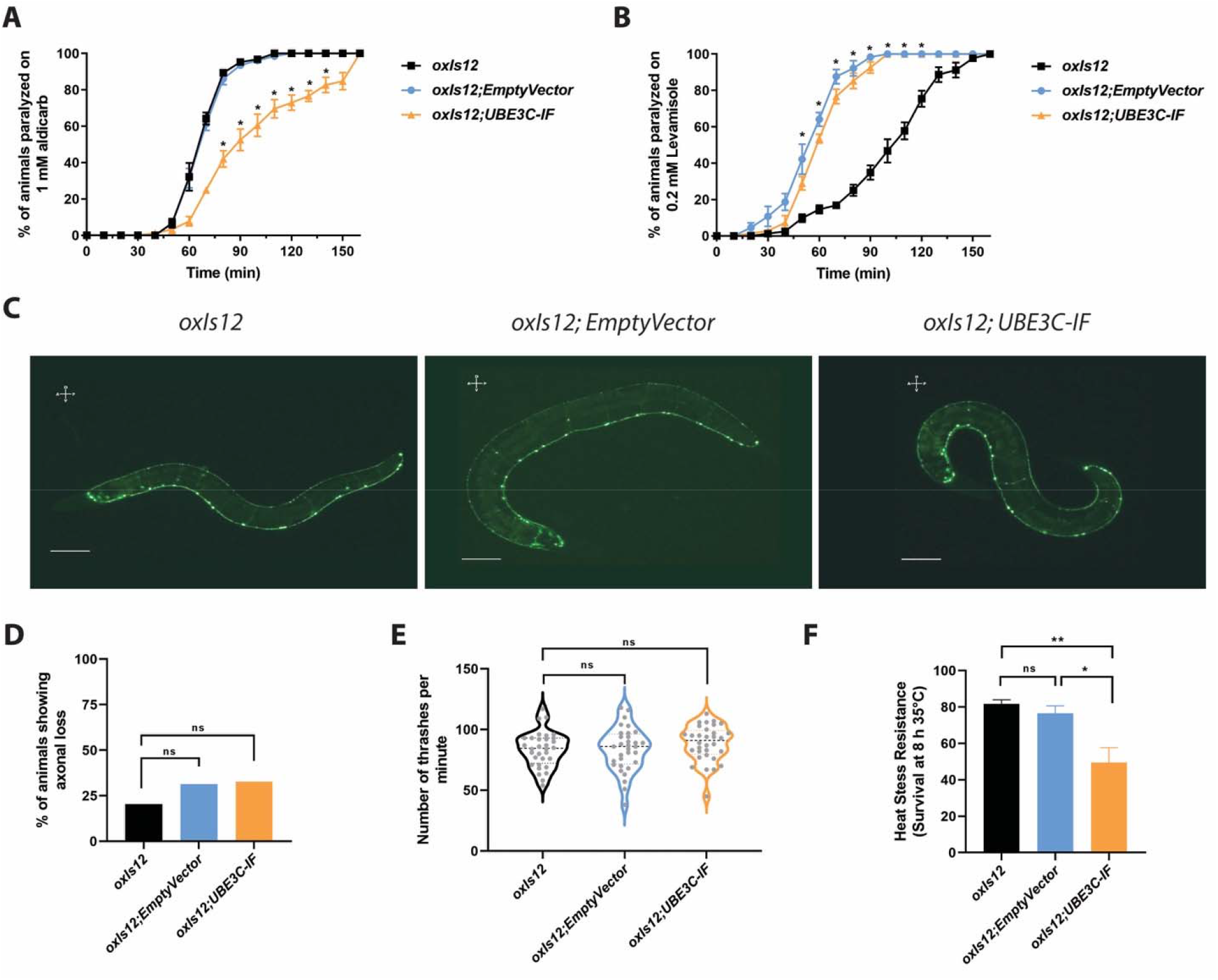
C. *elegans* transgenic animals overexpressing UBE3C-IF show synaptic transmission deficits and susceptibility to heat stress. (**A and B**) *C. elegans* strains were treated with 1 mM aldicarb and 0.2 mM Levamisole. In the presence of 1 mM aldicarb, *C. elegans* mutants carrying *UBE3C-IF* transgene showed significant resistance to aldicarb induced paralysis when compared to control animals suggesting synaptic transmission deficits. Upon treatment with levamisole, transgenic *C. elegans* strains showed increased paralysis when compared to *oxIs12* animals. However, there was no significant difference between transgenic strains carrying empty vector and *UBE3C-IF* transgene suggesting that the aldicarb phenotype observed in the *oxIs12;UBE3C-IF* animals are due to pre-synaptic deficits. 20 to 25 animals were used per biological replicate with a minimum of three biological replicates per strain. The mean ± SEM is presented. * p-value < 0.05, two-tailed unpaired t-test. **(C)** Representative images of day-1 old control (*oxIs12 and oxIs12; EmptyVector*) and mutant (*oxIs12; UBE3C-IF*) *C. elegans* strains. The ventral nerve cord (VNC) of the transgenic strains were observed to be normal and similar to the *oxIs12* animals with all the cell bodies and axons intact. Neurodegeneration was not observed in *oxIs12* animals overexpressing the *UBE3C-IF* transgene. Scale bar – 0.3 mm. **(D)** Quantification of axonal neurodegeneration. No significant axonal degeneration was identified in transgenic animals when compared to controls. Animals used for live imaging are as follows: *oxIs12* (n=91), *oxIs12;EmptyVector* (n=79) and *oxIs12;UBE3C-IF* (n=78). ns - not significant. **(E)** Quantification of body thrash assay. Analysis of the swimming (thrashing) behaviour of day 4 old *C. elegans* strains showed no locomotion deficits associated with *UBE3C-IF* overexpression when compared to controls. ns - not significant. **(F)** Heat stress resistance assay. *C. elegans* strains were exposed to a temperature of 35°C for 8 h and the survival percentage calculated following the heat exposure. Transgenic animals overexpressing the *UBE3C-IF* transgene showed a significant reduction in survival when compared to *oxIs12;EmptyVector* and *oxIs12* controls. Data was generated with 20 to 30 animals per genotype per replicate. A total of 4 experimental replicates were used for the heat stress resistance assay. Data is presented as mean ± SEM. One-way ANOVA and Tukey’s multiple correction test. * adjusted p-value = 0.01 and ** adjusted p-value = 0.003.

## Discussion

Distal hereditary motor neuropathies have benefited from next generation sequencing gene discovery and genetic diagnosis; however, a large proportion of cases (up to 50%) remain genetically unsolved. In such instances, there is a strong precedent for looking beyond the exome and querying the remainder of the genome for SV mutations. Several cases of SV – both typical and atypical (see Cutrupi *et al.*^29^ and Boyling *et al.*^84^ for a review) – have been reported to cause inherited peripheral neuropathies (IPN). Therefore, in the cases where protein-coding mutations cannot be identified, SV represents a possible mutational mechanism that may account for some of the genetically undiagnosed dHMN and other IPN cases^29^.

This study highlights the utility of this approach and represents the first available human neuronal model to examine the impact of the DHMN1 complex SV mutation. We have shown the 1.35 Mb complex insertion contains a partial duplication of the *UBE3C* gene which forms a novel gene-intergenic fusion (*UBE3C-IF*) by incorporating a pseudo-exon from intergenic sequence from within the DHMN1 locus. The *UBE3C-IF* forms a stable transcript that is not degraded by nonsense mediated decay and can readily be detected in mRNA from patient derived sMN. Chromosomal rearrangements such as somatic SVs are a common driver of fusion gene formation. Several publications have described this mechanism in cancer where gene fusions have been extensively studied.^85–92^ Typically, fusion genes involve two or more coding sequences (gene-gene fusions; e.g. *BCR-ABL*^85,86^ and *TMPRSS-ERG*^87^) resulting in chimeric mRNA transcripts that can form oncogenic “neo-antigens”.^93^ Fusions involving intergenic/non-coding sequences that result in cryptic (pseudo) exon formation are by comparison rare – only being reported by a few studies.^93–96^ Interestingly, the majority of genomic breakpoints in fusion genes are intergenic or intronic and are therefore not typically present in mRNA or protein-coding sequences^90,91,97^ making detection of gene-intergenic fusions difficult. This may account for the predominance of classical gene-gene fusions over gene-intergenic fusions in the literature to date. Thorough investigation of genetically unsolved and/or suspected SV cases using a combination of WGS and RNA-seq may therefore help to improve the diagnostic rate of gene-intergenic fusions in genetic disease and expand the spectrum of mutations causing inherited motor neuron disorders.

Studying the effects of these SV mutations is challenging as the size and complexity of the DNA rearrangement can increase the number of potential candidate causative genes. This makes interpretation of data from gene expression studies difficult particularly when alternative tissues are used in place of disease relevant tissue. In this study, we addressed this issue by generating a disease-relevant, tissue-specific model and applying rigorous, experimental-driven filtering to refine the list of probable disease-causing candidate genes. Our original hypothesis was that DHMN1 neuropathy is caused by transcriptional dysregulation. Using a positional cloning approach, we showed that using a targeted gene expression panel of 63 genes combined with Hi-C analysis was able to eliminate all but two high-priority candidate genes, *MNX1* and *UBE3C*. Western blot analysis showed no change in MNX1 and was therefore excluded as a causative candidate gene. The observed transcriptional upregulation of *MNX1* is likely a bystander effect of the SV mutation. MNX1 has been reported to autoregulate^98^ and this may explain why a corresponding difference was not observed at the protein level. In contrast, although *UBE3C* appeared to be upregulated in patient tissues, we showed that this was an artefact based on the primer/probe design detecting both *UBE3C-WT* and *UBE3C-IF* transcripts (Fig. 6D). The lack of meaningful gene dysregulation observed at the local transcriptional level was also observed globally with only 22 of 19,929 genes detected as differentially expressed. The absence of any discernible and meaningful aberrant transcriptional regulation provides further evidence against our original hypothesis and lends strong support for *UBE3C-IF* as the pathogenic candidate. This is further strengthened by the *UBE3C-IF* pseudo-exon using DNA sequences from within the previously mapped DHMN1 linkage region.^69^

The ubiquitin protein E3 ligases are a superfamily of over 600 genes responsible for the transfer of ubiquitin (Ub) to substrate proteins marked for degradation by the ubiquitin proteasome system (UPS^99,100^). UBE3C is a member of the HECT (homologous to E6-AP carboxyl terminus) class of E3 ligases which comprises 28 members.^101^ The activity of HECT E3 ligases is tightly regulated with respect to Ub linkage specificity and chain type, interaction with cognate E2 conjugating enzymes and substrate recognition.^102^ E3 ligases (in particular HECT E3 ligases) are reported to have important functions in neuronal development and migration, synaptic transmission^99^ and have been implicated in a variety of neurological and neurodevelopmental diseases including inherited peripheral neuropathy (see Ambrozkiewicz *et al.*,^99,103^ George *et al.,*^100^ and Lescouzeres *et al.*^104^ for review). We show that UBE3C is present in iPSC, MNP and sMN. This is not surprising given that the vast majority of E3 ligases (including *UBE3C*) are ubiquitously expressed^105^. Importantly however, these three tissues correspond to three distinct developmental stages suggesting that the *UBE3C* gene may be important in neuronal development. GO analysis of the 22 DEGs identified several enriched pathways related to nervous system development. However, given the lack of meaningful differential expression detected at both local and global transcriptional levels, it cannot be determined if this is related to UBE3C or an artefact of inducing neuronal differentiation. Furthermore, given that we show no observable change in *UBE3C-WT* mRNA expression, it may be unlikely that the term enrichment in DEGs is related to *UBE3C*.

E3 ligases are primary determinants of the substrate specificity of the UPS, however many of them are poorly characterised in this regard.^106^ The specificity of E3 ligase-substrate interactions contributes to the large diversity in the spatiotemporal control of ubiquitination.^106,107^ In order to preserve this, E3 ligases are tightly regulated by post-transcriptional modifications such as ubiquitination^108^ or by various homotypic and/or heterotypic interactions (reviewed in Balaji & Hoppe^109^). In this study, we observed that levels of full-length wild-type UBE3C are reduced in patient sMN harbouring the UBE3C-IF. Furthermore, we show overexpression of UBE3C-IF recapitulated the reduced endogenous UBE3C-WT levels in HeLa cells which was observed in patient sMN. Taken together, these data suggest that the presence of UBE3C-IF protein at higher concentrations affects UBE3C autoregulation resulting in reduced levels of endogenous UBE3C-WT protein. This is not unprecedented given that autoregulation has been observed among E3 ligase family members. For example, upon degradation of their respective substrates, the E3 ligases MDM2 and SIAH1 have been shown to self-regulate.^110,111^ This is triggered by increased cellular levels of these proteins following target substrate degradation. MDM2 and SIAH1 autoregulation begins with homodimerization proceeded by autoubiquitination resulting in proteasomal degradation.^110,111^ Recent evidence suggests that UBE3C can self-regulate through autoubiquitination^112^ and it is therefore possible that the observed reduction in UBE3C-WT in patient sMN is due to autoregulation triggered by the presence of UBE3C-IF. This might also explain the absence of the UBE3C-IF protein signature in sMN western blot. Aside from the aforementioned co-migration UBE3C-IF with a shorter tissue-specific isoform obscuring visualisation, it is possible that due to the rapid nature of UPS-mediated protein degradation, UBE3C-IF and UBE3C-WT are degraded simultaneously and too rapidly to observe UBE3C-IF without the use of proteasome inhibitors. This will be useful to investigate further in follow-up studies.

The maintenance of protein homeostasis is fundamental to the proper functioning of cells. To achieve this, cells must balance protein synthesis with protein degradation.^113^ The transfer of Ub to targeted substrates is fundamental to this process. Mutation or dysfunction of genes within the system results in the accumulation of ubiquitinated inclusions which have been associated with several neurodegenerative diseases^114,115^ including amyotrophic lateral sclerosis (ALS)^116–119^. Furthermore, perturbed protein homeostasis is implicated in ALS. UBE3C associates with the proteasome and assembles Lys-29 and Lys-48 linked polyUb chains.^113,120–122^ Knockdown experiments in yeast have shown that the absence of *Hul5p* (orthologous to human *UBE3C*) results in a reduction in the processivity of the proteasome making it less able to degrade stable proteins leaving behind partially degraded remnants.^123,124^ Recent studies have expanded on these findings and have shown the UBE3C protects against accumulation of partially degraded substrates arising due to partial proteolysis.^113^

To investigate the role of the *UBE3C* gene-intergenic fusion on neuron morphology and nervous system function, we generated transgenic *C. elegans* that either overexpressed *UBE3C-IF* transcript or the empty vector backbone in animals that carry the transgene *oxIs12*[*unc-47::GFP*]. Defects associated with the neuromuscular junction (NMJ) are reported for peripheral neuropathies.^125^ UBE3C-IF animals displayed a resistance phenotype in the presence of 1 mM aldicarb (Fig. 8A). Aldicarb resistance is observed in mutants with less acetylcholine in the synaptic cleft, suggesting potential deficits in neurotransmitter release in UBE3C-IF animals. In *C. elegans*, overexpression of UBE3C-IF did not affect neuron morphology and animal locomotion (Fig. 8C-E). Upon exposure to heat stress, UBE3C-IF animals showed reduced survival suggesting a defective response to temperature-induced protein unfolding stress (Fig 8F). Interestingly, knockdown of Hulp5 (the yeast ortholog of *UBE3C*) was found to compromise the recovery of yeast after heat shock.^83^ Similarly, knockout of UBE3C in human cells renders them more susceptible to heat shock following treatment with a heat shock protein 90 (hsp90) inhibitor.^113^ Taken together, our DHMN1 in vivo model shows that the UBE3C-IF causes synaptic transmission deficits and defective stress response resulting in reduced animal survival.

## Conclusions

This study has presented the first patient derived sMN model for DHMN1. Moreover, this study is the first report of a gene-intergenic fusion (*UBE3C-IF*) causing a motor neuron disorder and has therefore expanded the spectrum of mutations known to cause motor neuron diseases. This study has shed light on a new disease mechanism in motor neuron diseases and underscores the importance of looking beyond the exome when studying inherited diseases with unknown genetic aetiology. Furthermore, we show the potential of *UBE3C-IF* using an *in vivo* model to elucidate pathogenic processes through highlighting synaptic dysfunction and compromised proteasome processivity as plausible disease processes underlying DHMN1 pathogenesis. Further follow-up is needed to define the precise mechanism of action by which UBE3C-IF and UBE3C-WT interact and to define the targets of UBE3C-WT. This work demonstrates the novelty of gene-intergenic fusions as an important and understudied mechanism for motor neuron disease that will provide essential new knowledge and inform avenues for treatment and therapies.

## Author contributions

MLK and ANC devised concept and study design. ANC, RN and GPS devised the experiments. ANC performed the cellular experiments and analysed the data. GPS performed and analysed the western blot experiments. RN engineered the transgenic *C. elegans* model and performed and analysed *C. elegans* experiments. BN advised on *C. elegans* experimental design. BG performed cloning, construct production and *in silico* splice analysis. KL provided bioinformatic support for mapping and analysis of Hi-C data. AB provided assistance in the culturing, maintenance and differentiation of iPSC and motor neurons and RNA extractions. ME assisted in the preparation of patient fibroblasts samples for iPSC reprograming. DM and MU provided SN38-P for use in differentiation protocols. MAS trained ANC in the use of iPSC and motor neuron differentiation. RCYL advised on NanoString and mRNA-seq experimental design. GAN contributed patients to the study. ANC prepared the manuscript. MLK, RN, GPS, BG, KL, AB, SV, RCYL, GAN and MAS provided advice and assistance in the preparation of manuscript.

## Acknowledgements

The authors thank the CMT families who have participated in the research and volunteered skin biopsies for the generation of iPSC under informed consent. We also acknowledge and thank Dr Renata Maciel for her invaluable technical assistance with iPSC and sMN work presented in this study. Some *C. elegans* strains were obtained from the Caenorhabditis Genetics Center (CGC) funded by NIH Office of Research Infrastructure programs (P40 OD010440). We would like to thank Paul Davis and Prof. Tim Schedl for assistance with obtaining WormBase strain accession numbers. Strains generated in this study can be obtained by contacting the corresponding author.

## Supplementary material

Supplementary material is available at *Brain* online.

