## Supplementary Figures for "Novel gene-intergenic fusion involving ubiquitin E3 ligase UBE3C causes distal hereditary motor neuropathy: A new mechanism for motor neuron degeneration"

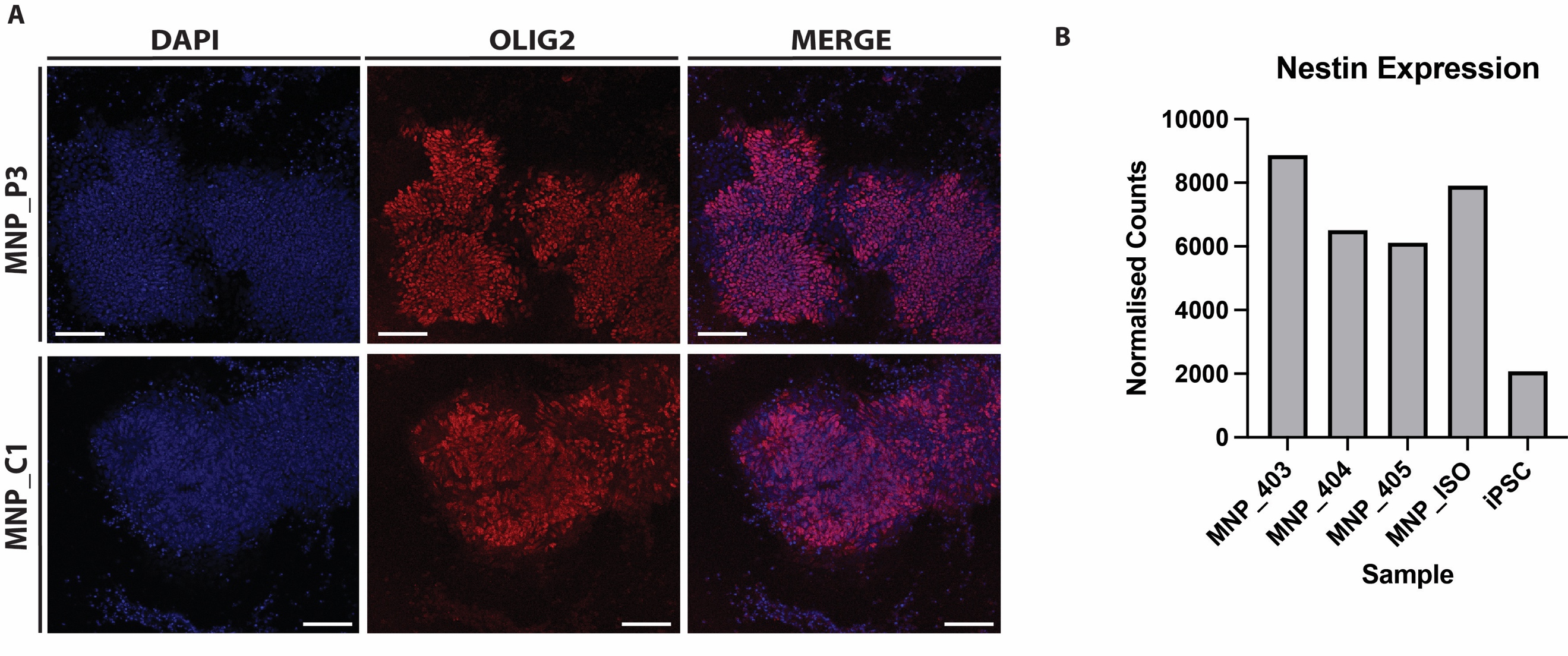

**Supplementary Figure S1: MNPs derived from patient and control iPSCs express markers of motor neuron progenitors. (A)** Immunofluorescence confirms identity of MNPs by expression of OLIG2 (red) within the nuclei (blue) of progenitor cells. Scale bar 90 µm. **(B)** Expression levels of MNP/NSC marker *NES* (Nestin) in iPSC and MNP.

**RNA-seq diagnostics and additional analyses.**

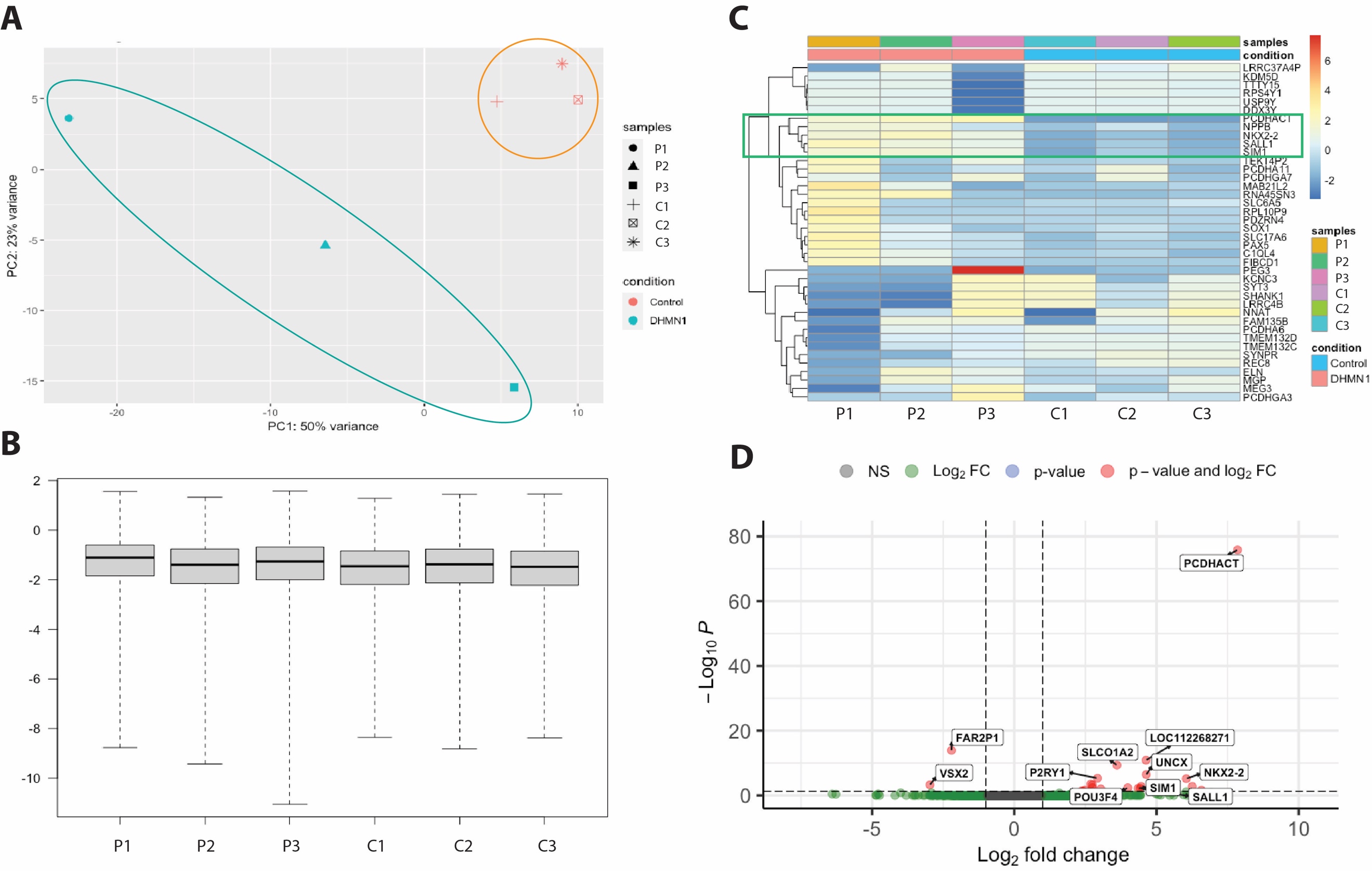
Principal component analysis (PCA) on regularised log (rlog) transformed data demonstrated the DHMN1 samples clustered separately from the controls (Supplementary Figure S2A). Analysis of Cook’s differences found no sample outliers (Supplementary Figure S2B). Gene clustering analysis using a subset of the top 40 most variable genes in the dataset following rlog transformation indicated minimal covariance between conditions (Supplementary Figure S2C). This was recapitulated with a subset of the top 100 most variable genes (Supplementary Figure S3B). Similarly, when genes were ranked by row mean, no covariance was observed between conditions when clustering was performed on a subset of the top 100 genes (supplementary Figure S3A).

**Supplementary Figure S2: Exploratory plots of RNA-seq data reveal no evidence of biologically relevant differential expression (A)** Principal component analysis (PCA) plot of DHMN1 and control sMN show distinct clustering by affection status **(B)** Box plot of Cook’s distances indicate absence of sample outliers **(C)** Hierarchical gene clustering analysis of the top 40 most variable genes shows minimal covariance between patient and control sMN. Each row corresponds to a gene and each column corresponds to a DHMN1 or control sMN sample. The colour of each rectangle indicates the amount by which gene counts for a given gene in a specific sample deviates from the average expression of that gene across all samples. Green box indicates a block of genes that covaries between conditions. Data is rlog transformed. **(D)** Volcano plot of expressed genes. Green points represent genes that pass the log2FC threshold but do not reach significance. Grey points represent genes that do not pass significance or log2FC thresholds. Red points represent differentially expressed features. The shape of the volcano plot is flat indicating minimal biologically relevant differential expression between DHMN1 and control sMN.

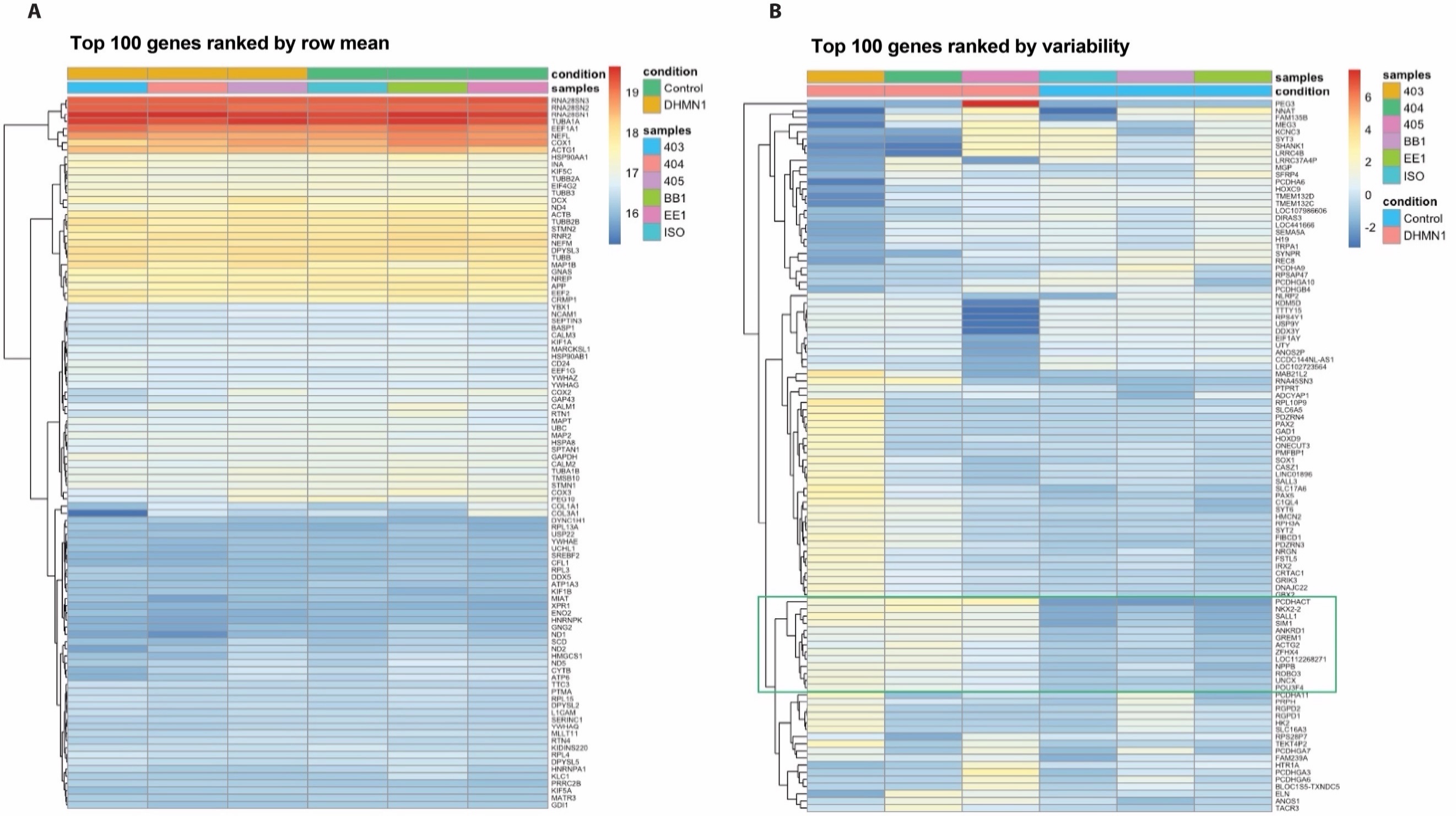

**Supplementary Figure S3: Hierarchical gene clustering analysis reveals minimal covariance between DHMN1 and control sMN. (A)** Top 100 genes ranked by row mean and clustered. Each row corresponds to a gene and each column corresponds to a DHMN1 or control sMN sample. The colour of each rectangle indicates the rlog transformed count value of genes in a sample. Clustering reveals no covariance of genes between samples. **(B)** Top 100 genes ranked by variability and clustered. Each row corresponds to a gene and each column corresponds to a DHMN1 or control sMN sample. The colour of each rectangle indicates the rlog transformed count value of genes in a sample. Clustering reveals no covariance of genes between samples. The colour of each rectangle indicates the amount by which a gene’s counts in a specific sample deviates from the gene’s average across all samples. Green box indicates a block of genes that covaries between conditions. Data is rlog transformed.

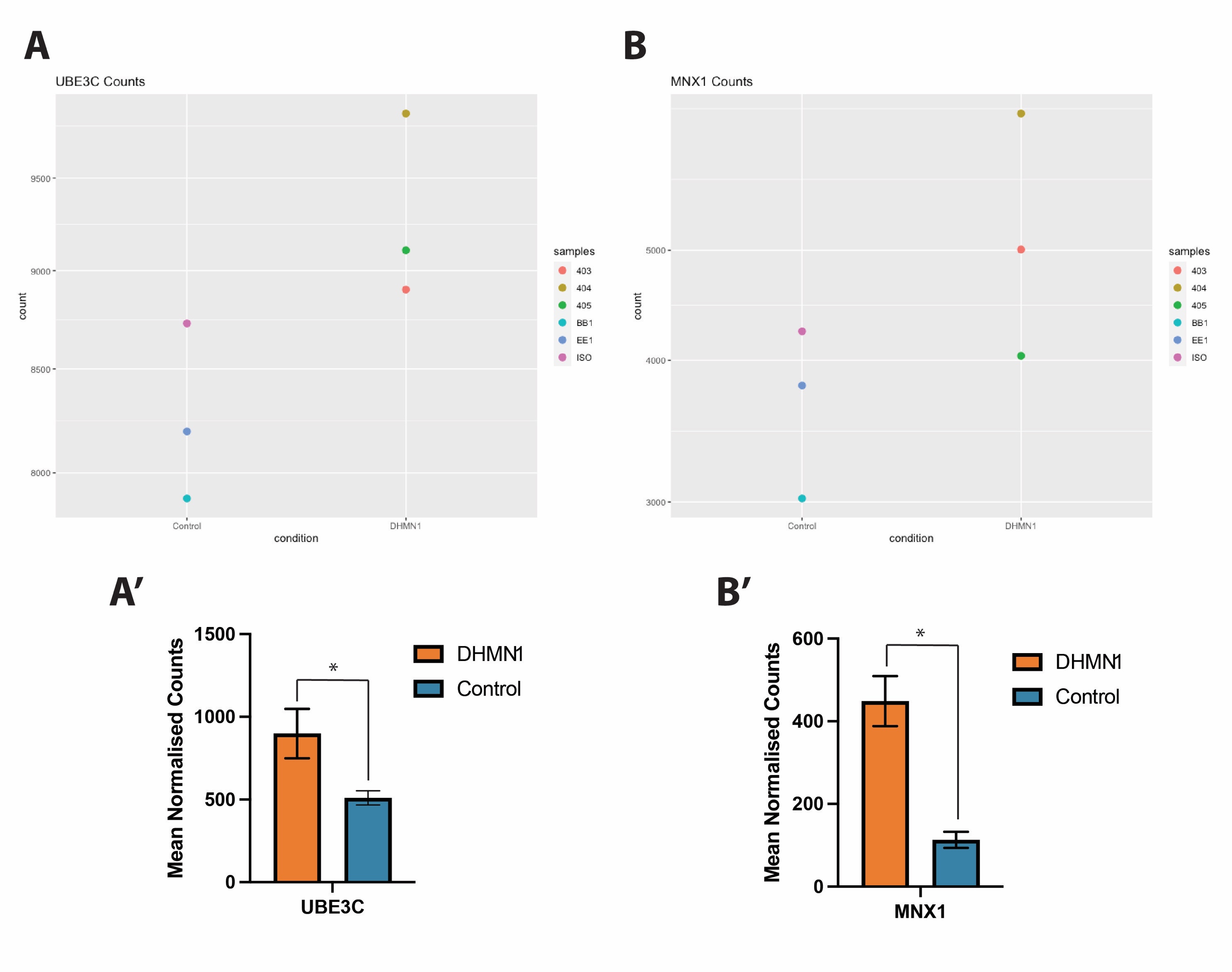

**Supplementary Figure S4: Normalised RNA-seq read counts for genes identified as differentially expressed in nanostring.** Whilst expression of *UBE3C* and *MNX1* from RNA-seq data display the same pattern of upregulation (**A** and **B** respectively) as observed with the nanostring assay **(A’-B’)**, this upregulation did not reach significance.

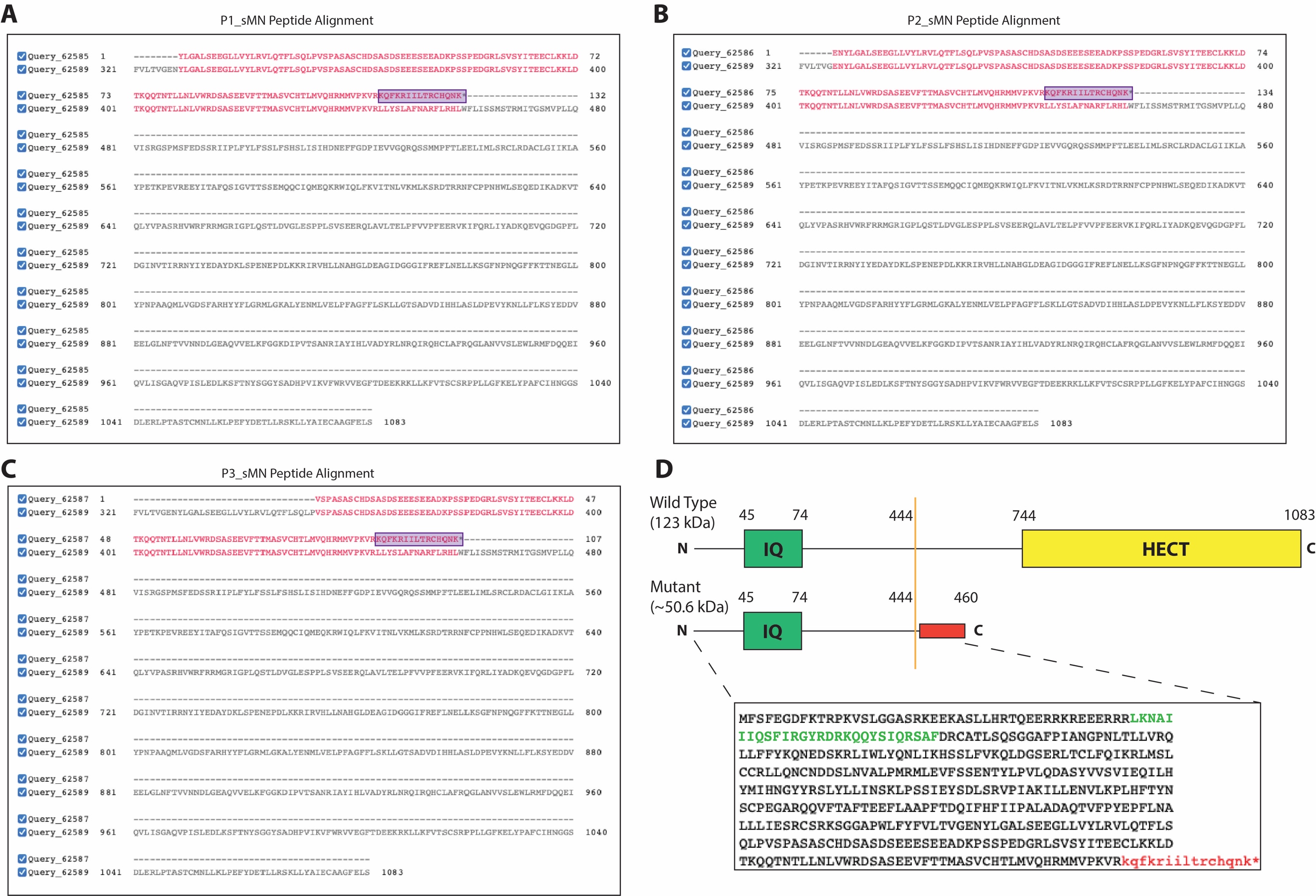

**Supplementary Figure S5: Peptide sequence alignments for UBE3C-IF. (A-C)** Predicted peptide sequences of the exon 10/pseudo-exon junction obtained from Arriba fusion gene detection algorithm were aligned using BLASTP multisequence alignment. Red letters indicate amino acids that were aligned. Purple boxes indicate amino acids from the predicted pseudo-exon that do not align to UBE3C wild-type protein (NP_055486.2). **(D)** Schematic representation of the structure of wild-type UBE3C and UBE3C-IF proteins with the expanded full predicted peptide sequence of UBE3C-IF shown below. The novel UBE3C-IF transcript shares the first 444 aa of wild-type UBE3C. Coloured letters correspond to annotated protein domains. IQ: Calmodulin binding domain (green); HECT: Homologous to E6AP Carboxyl Terminus domain (yellow). Red box corresponds to pseudo-exon amino acids. Solid orange line indicates breakpoint.

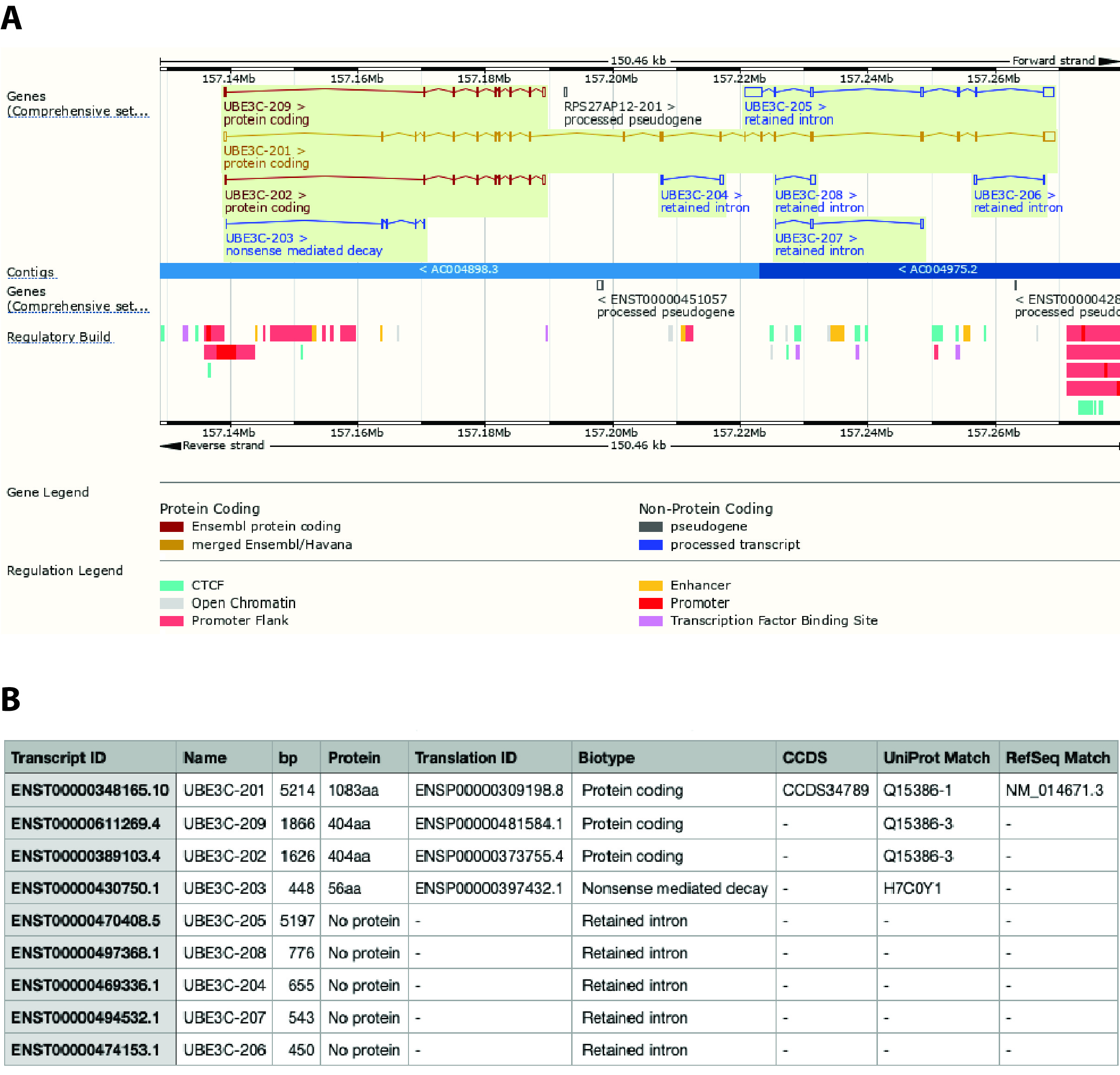

**Supplementary Figure S6: Peptide sequence alignments for UBE3C-IF. (A)** Graphical display of annotated UBE3C transcripts and protein isoforms. UBE3C-201 corresponds to RefSeq ID: NP_055486.2. **(B)** Table of information pertaining to transcripts and isoforms in (A). All data was obtained from the Ensembl database (<https://asia.ensembl.org/Homo_sapiens/Gene/Summary?db=core;g=ENSG00000009335;r=7:157138916-157269370>).

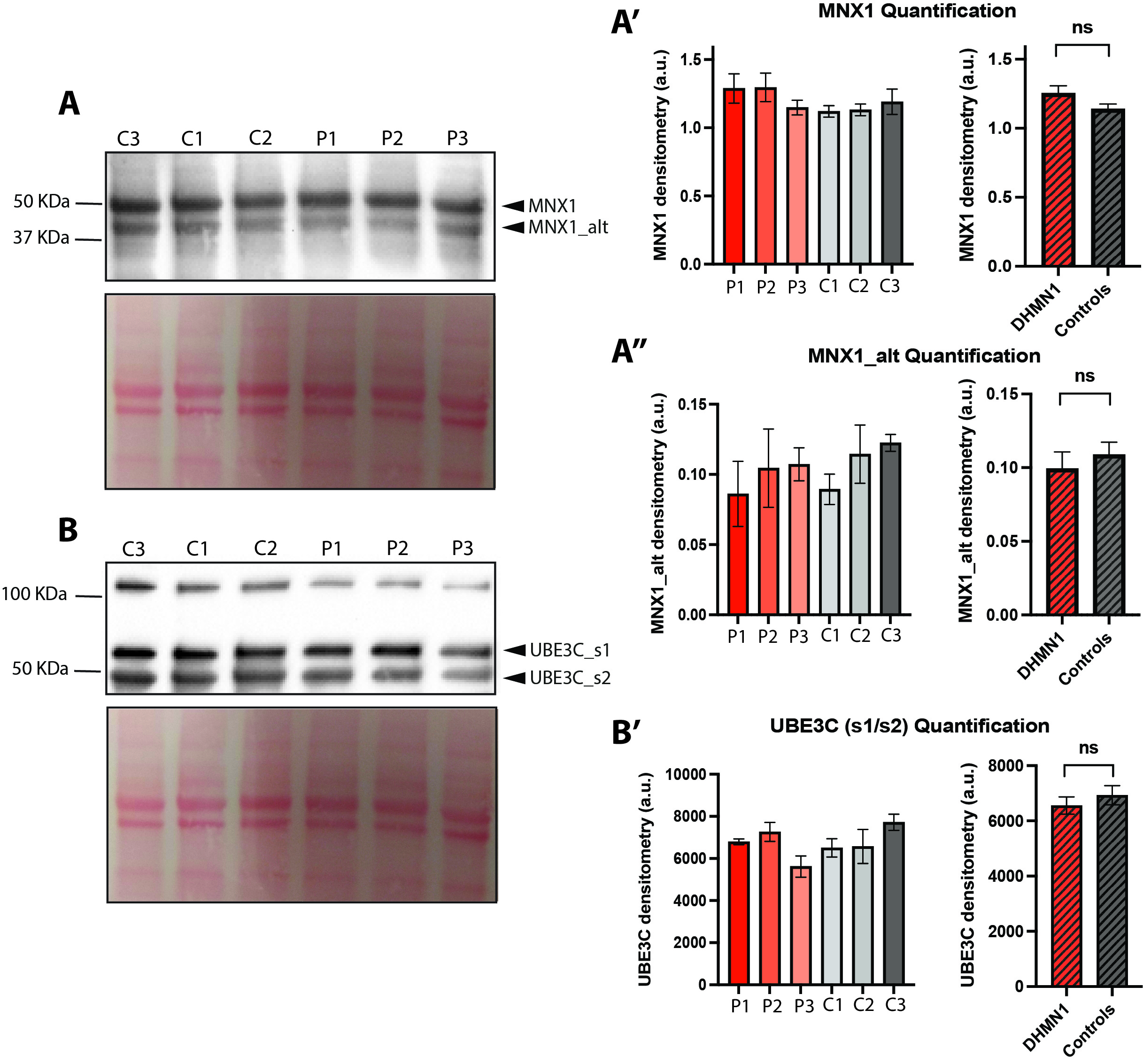

**Supplementary Figure S7: Quantifications of MNX1 and UBE3C alternate isoforms.** Representative western blot of **(A)** MNX1 (top panel) and **(B)** UBE3C (top panel). Ponceau stain is shown below (bottom panels). **(A’-A’’)** Quantification of MNX1 (top band) and MNX1_alt (bottom band) levels are not different in DHMN1 sMN protein lysates when compared to controls. **(B’)** Quantification of the probable short isoforms UBE3C_s1/s2 show no difference in DHMN1 sMN protein lysates when compared to controls.

**Original blots and gels.**

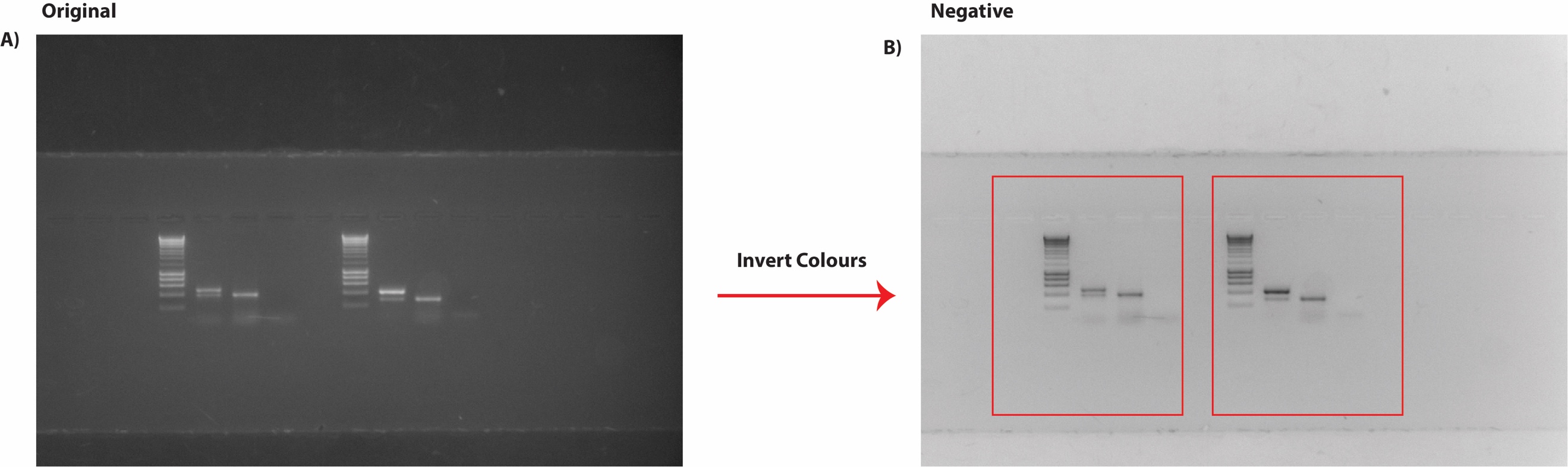

**Supplementary Figure S8: Processing of gel electrophoresis data for representation in Figure 1. (A)** Original photograph of gel. **(B)** A negative colour image of the gel was created using Adobe Illustrator via the ‘Invert Colours’ function. Cropping of the photograph is indicated by red boxes.

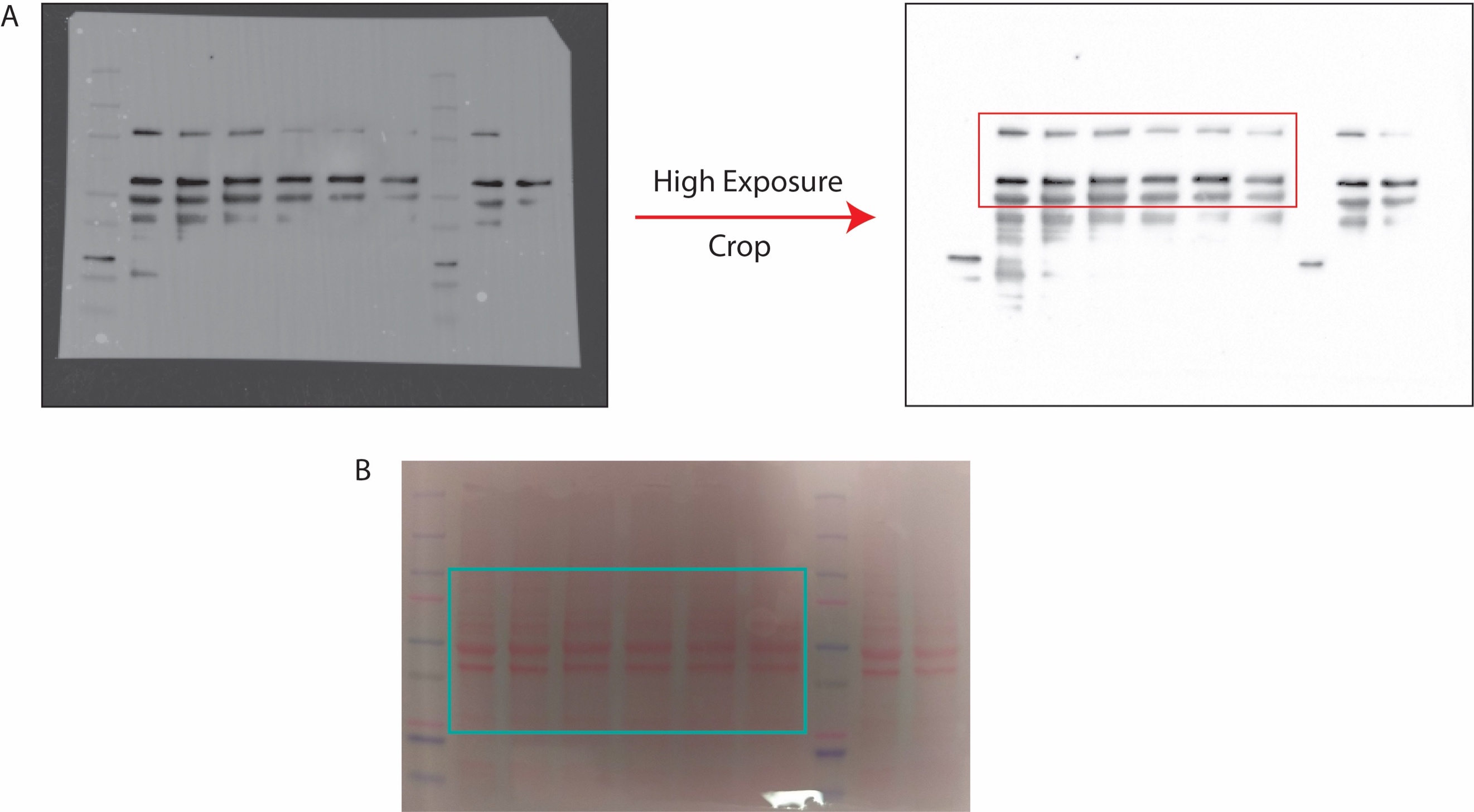

**Supplementary Figure S9: Processing of western blot images for representation in Figure 7A.** **(A)** Standard (left) and high exposure (right) photographs of western blot from sMN cell lysates. Images were cropped (red boxes) using Adobe Illustrator. **(C)** Ponceau stain. Region used for normalisation is indicated by the green box and is represented in figure 7B. Cropping was performed in Adobe Illustrator.

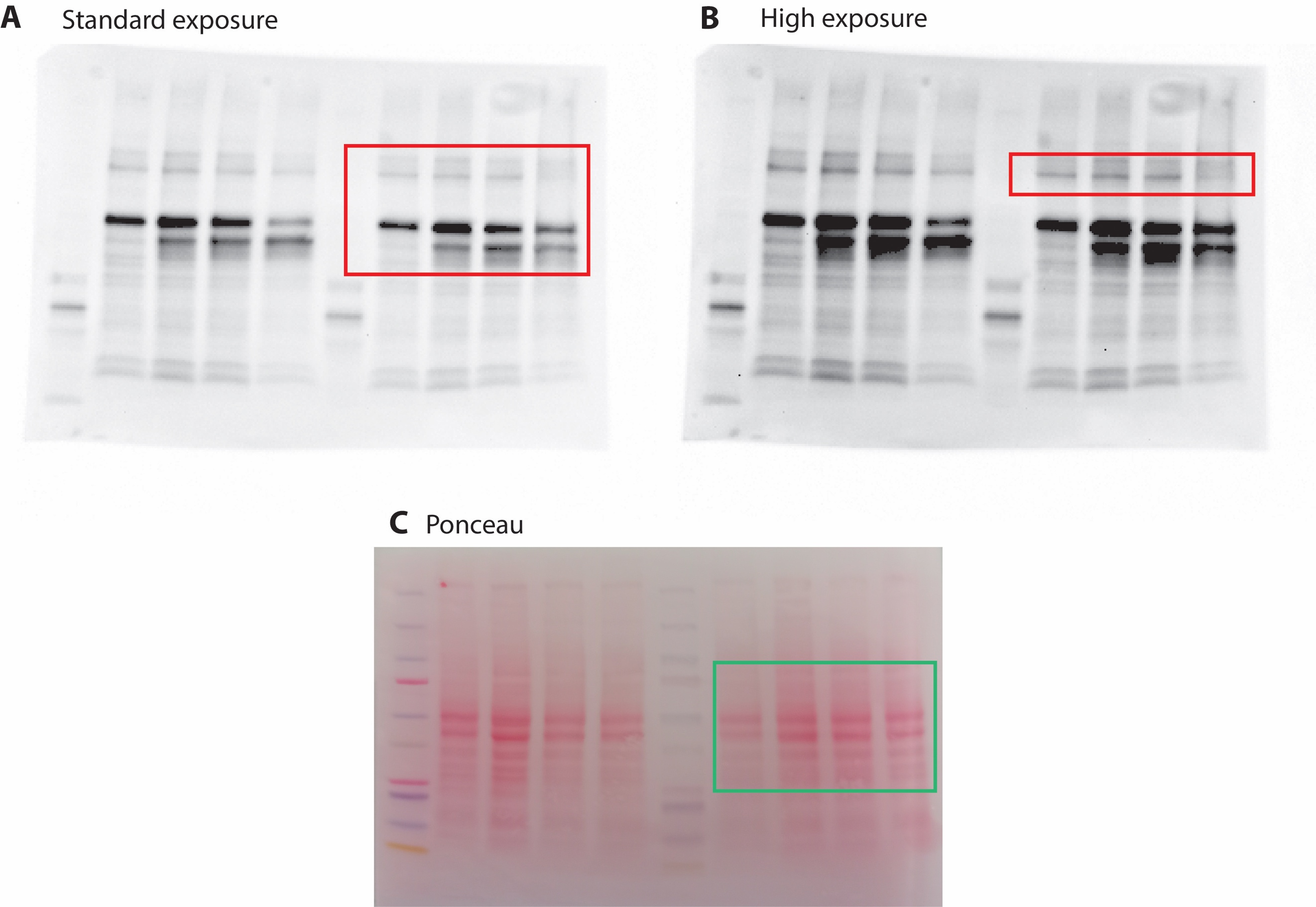

**Supplementary Figure S9: Processing of western blot images for representation in Figure 7B.** **(A)** Standard and **(B)** high exposure photographs of western blot from HeLa cell lysates. Images were cropped (red boxes) using Adobe Illustrator. **(C)** Ponceau stain. Region used for normalisation is indicated by the green box and is represented in figure 7B. Cropping was performed in Adobe Illustrator.
