## Supplementary Tables for "Novel gene-intergenic fusion involving ubiquitin E3 ligase UBE3C causes distal hereditary motor neuropathy: A new mechanism for motor neuron degeneration"

**Supplementary Table S1: TaqMan probes used for *UBE3C-IF* assay.**

TG: Target gene, HKG: Housekeeping gene

| **Gene** | **ID** | **Company** | **Exon Boundary** | **Type** |
| --- | --- | --- | --- | --- |
| *UBE3C* | Hs00904532_m1 | Applied Biosystems | 2-3 | TG |
| *UBE3C* | Hs00904531_m1 | Applied Biosystems | 22-23 | TG |
| *GAPDH* | Hs99999905_m1 | Applied Biosystems | 2-2 | HKG |
| *RPLPO* | Hs99999902_m1 | Applied Biosystems | 3-3 | HKG |
| *RPL30* | Hs00265497_m1 | Applied Biosystems | 4-5 | HKG |

**Supplementary Table S2: Antibodies used in molecular analyses**

IF: Immunofluorescence, WB: Western Blot

| **Antigen** | **Host** | **Company** | **Catalogue #** | **Concentration** | **Use** |
| --- | --- | --- | --- | --- | --- |
| MNX1 | Mouse | Sigma | HPA071717 | 1:500 | IF |
|  |  |  |  | 1:500 | WB |
| OCT4 | Rabbit | Cell Signalling | 2840 | 1:400 | IF |
| SOX2 | Rabbit | Cell Signalling | 3579 | 1:400 | IF |
| NANOG | Rabbit | Cell Signalling | 4903 | 1:400 | IF |
| OLIG2 (clone 211F1.1) | Mouse | Millipore | MABN50 | 1:100 | IF |
| NF68 | Chicken | Abcam | Ab24520 | 1:1000 | IF |
| TUBB3 | Mouse | Sigma | T2220 | 1:1000 | IF |
| UBE3C | Rabbit | Invitrogen | PA5-110540 | 1:100 | IF |
|  |  |  |  | 1:500 | WB |

**Supplementary Table S3: Targeted Nanostring panel for genes within a 6 Mb interval flanking the DHMN1 locus**

proximal: towards centromere, distal: towards telomere

| **Gene Name** | **Location** | **Distance From Insertion Site** | **Class** |
| --- | --- | --- | --- |
| *ABCB8* | Flank | 2.5 Mb - 3 Mb proximal | Target |
| *ABCF2* | Flank | 2 Mb - 2.5 Mb proximal | Target |
| *ACTR3B* | Flank | < 1 Mb proximal | Target |
| *AGAP3* | Flank | 2.5 Mb - 3 Mb proximal | Target |
| *AOC1* | Flank | 2.5 Mb - 3 Mb proximal | Target |
| *ASB10* | Flank | 2 Mb - 2.5 Mb proximal | Target |
| *ASIC3* | Flank | 2.5 Mb - 3 Mb proximal | Target |
| *ATG9B* | Flank | 2.5 Mb - 3 Mb proximal | Target |
| *BLACE* | Flank | 1.5 Mb - 2 Mb distal | Target |
| *CDK5* | Flank | 2.5 Mb - 3 Mb proximal | Target |
| *CHPF2* | Flank | 2 Mb - 2.5 Mb proximal | Target |
| *CNPY1* | Flank | 1.5 Mb - 2 Mb distal | Target |
| *CRYGN* | Flank | 2 Mb - 2.5 Mb proximal | Target |
| *DNAJB6* | Flank | > 3 Mb distal | Target |
| *DPP6* | Flank | < 500 kb distal | Target |
| *EN2* | Flank | 1.5 Mb - 2 Mb distal | Target |
| *FABP5P3* | Flank | 1.5 Mb - 2 Mb proximal | Target |
| *FASTK* | Flank | 2.5 Mb - 3 Mb proximal | Target |
| *GALNT11* | Flank | 1.5 Mb - 2 Mb proximal | Target |
| *GALNTL5* | Flank | 1.5 Mb - 2 Mb proximal | Target |
| *GAPDH* | N/A | N/A | Housekeeping |
| *GBX1* | Flank | 2 Mb - 2.5 Mb proximal | Target |
| *GIMAP1* | Flank | 2.5 Mb - 3 Mb proximal | Target |
| *GIMAP2* | Flank | 2.5 Mb - 3 Mb proximal | Target |
| *GIMAP3* | Flank | 2.5 Mb - 3 Mb proximal | Target |
| *GIMAP4* | Flank | 2.5 Mb - 3 Mb proximal | Target |
| *GIMAP5* | Flank | 2.5 Mb - 3 Mb proximal | Target |
| *HTR5A* | Flank | 1 Mb - 1.5 Mb distal | Target |
| *HTR5A-AS1* | Flank | 1 Mb - 1.5 Mb distal | Target |
| *INSIG1* | Flank | 1.5 Mb - 2 Mb distal | Target |
| *IQCA1L* | Flank | 2 Mb - 2.5 Mb proximal | Target |
| *KCNH2* | Flank | 2.5 Mb - 3 Mb proximal | Target |
| *KMT2C* | Flank | 1.5 Mb - 2 Mb proximal | Target |
| *LINC00244* | Insertion | Insertion | Target |
| *LINC01006* | Insertion | Insertion | Target |
| *LINC01287* | Flank | < 500 kb proximal | Target |
| *LMBR1* | Insertion | Insertion | Target |
| *LOC101929998* | Flank | < 500 kb distal | Target |
| *LOC389602* | Insertion | Insertion | Target |
| *MNX1* | Insertion | Insertion | Target |
| *MNX1-AS1* | Insertion | Insertion | Target |
| *MNX1-AS2* | Insertion | Insertion | Target |
| *MRPL19* | N/A | N/A | Housekeeping |
| *NOM1* | Insertion | Insertion | Target |
| *NOS3* | Flank | 2.5 Mb -3 Mb proximal | Target |
| *NUB1* | Flank | 2 Mb – 2.5 Mb proximal | Target |
| *PAXIP1* | Flank | 1 Mb – 1.5 Mb distal | Target |
| *PAXIP1-AS1* | Flank | 1 Mb – 1.5 Mb distal | Target |
| *PAXIP1-AS2* | Flank | 1 Mb – 1.5 Mb distal | Target |
| *PP1A* | N/A | N/A | Housekeeping |
| *PRKAG2* | Flank | 2 Mb – 2.5 Mb proximal | Target |
| *PRKAG2-AS1* | Flank | 2 Mb – 2.5 Mb proximal | Target |
| *PTPRN2* | Flank | > 3 Mb distal | Target |
| *RBM33* | Flank | 2 Mb - 2.5 Mb distal | Target |
| *RHEB* | Flank | 2 Mb - 2.5 Mb proximal | Target |
| *RNF32* | Insertion | Insertion | Target |
| *RPLPO* | N/A | N/A | Housekeeping |
| *SHH* | Flank | 2 Mb - 2.5 Mb distal | Target |
| *SLC4A2* | Flank | 2.5 Mb - 3 Mb proximal | Target |
| *SMARCD3* | Flank | 2 Mb – 2.5 Mb proximal | Target |
| *TBP* | N/A | N/A | Housekeeping |
| *TMEM176A* | Flank | 2.5 Mb - 3 Mb proximal | Target |
| *TMEM176B* | Flank | 2.5 Mb - 3 Mb proximal | Target |
| *TMUB1* | Flank | 2.5 Mb - 3 Mb proximal | Target |
| *UBE3C* | Insertion (partial) | Insertion | Target |
| *WDR86* | Flank | 2-2.5 Mb proximal | Target |
| *WDR86-AS1* | Flank | 2-2.5 Mb proximal | Target |
| *XRCC2* | Flank | < 1 Mb proximal | Target |

**Supplementary Table S4: Dysregulated genes identified in DHMN1 MNP and iPSC using a custom Nanostring nCounter Assay**

| **MNP Dysregulated Genes** | | | | | |
| --- | --- | --- | --- | --- | --- |
| **Target** | **dHMN1 Mean** | **Reference (Ctrl)** | **FC** | **Change in dHMN1** | **DE^*^** |
| *CHPF2* | 231.03 | 147.49 | 1.57 | up-regulated | Yes |
| *INSIG1* | 1186.27 | 826.53 | 1.44 | up-regulated | Yes |
| *LINC01006* | 218.27 | 123.55 | 1.77 | up-regulated | Yes |
| *LMBR1* | 2217.58 | 1437.57 | 1.54 | up-regulated | Yes |
| *MNX1* | 86.01 | 30.65 | 2.81 | up-regulated | Yes |
| *NOM1* | 355.46 | 249.01 | 1.43 | up-regulated | Yes |
| *PRKAG2* | 472.05 | 79.56 | -1.4 | down-regulated | Yes |
| *TMEM176A* | 668.34 | 1222.08 | -1.83 | down-regulated | Yes |
| *TMEM176B* | 631.51 | 1621.46 | -2.57 | down-regulated | Yes |
| *UBE3C* | 1907.85 | 920.39 | 2.07 | up-regulated | Yes |
| *XRCC2* | 312.11 | 117.51 | 1.48 | up-regulated | Yes |
| **iPSC Dysregulated Genes** | | | | | |
| **Target** | **dHMN1 Mean** | **Reference (Ctrl)** | **FC** | **Change in dHMN1** | **DE^*^** |
| *ABCB8* | 249.68 | 149.67 | 1.67 | up-regulated | Yes |
| *AGAP3* | 2403.84 | 1369.52 | 1.76 | up-regulated | Yes |
| *GALNT11* | 1350.61 | 190.65 | 1.33 | up-regulated | Yes |
| *LINC01006* | 533.04 | 205.53 | 2.59 | up-regulated | Yes |
| *LMBR1* | 3068.91 | 1628.76 | 1.88 | up-regulated | Yes |
| *NOM1* | 1148.38 | 647.58 | 1.77 | up-regulated | Yes |
| *SLC4A2* | 1539.92 | 997.71 | 1.54 | up-regulated | Yes |
| *SMARCD3* | 373.28 | 139.43 | 2.68 | up-regulated | Yes |
| *TMUB1* | 999.8 | 733.30 | 1.36 | up-regulated | Yes |
| *UBE3C* | 4038.31 | 1760.96 | 2.29 | up-regulated | Yes |
| *WDR86* | 864.97 | 648.61 | 1.33 | up-regulated | Yes |
| *XRCC2* | 1238.51 | 928.51 | 1.33 | up-regulated | Yes |

* DE call is determined based on the *DE call* test developed by Nanostring for nCounter data without replicates. It determines whether observed count differences between individual samples are larger than can be explained by technical noise alone (MAN-C0011-04)^1^. No *p-value* output is obtained using this test.

**Supplementary Table S5: Dysregulated genes identified in DHMN1 sMN using a custom Nanostring nCounter Assay**

| **Target** | **dHMN1 Mean** | **Ctrl Mean** | **LogFC** | **Change in dHMN1** | ***p*- value** |
| --- | --- | --- | --- | --- | --- |
| *MNX1* | 448.79 | 113.07 | 3.97 | up-regulated | 0.01386124 |
| *UBE3C* | 898.95 | 508.79 | 1.77 | up-regulated | 0.01414661 |
| *TMEM176B* | 35.98 | 183.49 | -5.1 | down-regulated | 0.02799794 |
| *SHH* | 193.32 | 49.36 | 3.92 | up-regulated | 0.03211705 |

**Supplementary Table S6: Quality Control and Mapping Statistics for RNA-seq Data**

| **Raw Data Statistics** | | | | | | | | | | |
| --- | --- | --- | --- | --- | --- | --- | --- | --- | --- | --- |
| **Batch** | **Sample id** | | | **Total read bases** | | **Total reads** | **GC (%)** | | **Q20 (%)** | **Q30 (%)** |
| N/A | P1 | | | 16,640,220,234 | | 110,200,134 | 49.99 | | 98.33 | 95.07 |
| N/A | P2 | | | 18,166,476,592 | | 120,307,792 | 50.08 | | 98.44 | 95.32 |
| N/A | P3 | | | 19,929,301,360 | | 131,982,130 | 49.75 | | 98.36 | 95.12 |
| N/A | C1 | | | 19,828,221,324 | | 131,312,724 | 49.94 | | 98.24 | 94.93 |
| N/A | C2 | | | 18,281,583,590 | | 121,070,090 | 49.79 | | 98.28 | 94.97 |
| N/A | C3 | | | 19,885,884,298 | | 131,694,598 | 49.60 | | 98.25 | 94.90 |
| **Average Total Raw Reads** | | | | | | 124,427,911 | | | | |
| **Trimmed Data Statistics** | | | | | | | | | | |
| **Batch** | **Sample id** | | | **Total read bases*** | | **Total reads** | **GC (%)** | | **Q20 (%)** | **Q30 (%)** |
| N/A | P1 | | | 16,036,353,394 | | 109,126,924 | 49.97 | | 98.86 | 95.84 |
| N/A | P2 | | | 17,546,053,268 | | 119,232,914 | 50.06 | | 98.94 | 96.04 |
| N/A | P3 | | | 19,181,722,408 | | 130,766,072 | 49.72 | | 98.88 | 95.89 |
| N/A | C1 | | | 18,966,151,086 | | 129,349,620 | 49.66 | | 98.87 | 95.84 |
| N/A | C2 | | | 17,534,209,115 | | 119,826,344 | 49.75 | | 98.85 | 95.79 |
| N/A | C3 | | | 19,077,876,355 | | 130,283,856 | 49.58 | | 98.83 | 95.74 |
| **Average Total Trimmed Reads** | | | | | | 123,097,622 | | | | |
| **Raw Data Mapping Statistics** | | | | | | | | | | |
| **Batch** | | **Sample id** | **# processed reads** | | **# mapped reads (%)** | | | **# unmapped reads (%)** | | |
| N/A | | P1 | 109,126,924 | | 107,949,591  (98.92%) | | | 1,177,333  (1.08%) | | |
| N/A | | P2 | 119,232,914 | | 178,870,341  (98.86%) | | | 1,362,573  (1.14%) | | |
| N/A | | P3 | 130,766,072 | | 129,347,512  (98.92%) | | | 1,418,560  (1.08%) | | |
| N/A | | C1 | 129,723,764 | | 128,332,988  (98.83%) | | | 1,390,776  (1.07%) | | |
| N/A | | C2 | 119,826,344 | | 118,293,731  (98.72%) | | | 1,532,613  (1.28%) | | |
| N/A | | C3 | 130,283,856 | | 128,832,736  (98.89%) | | | 1,415,120  (1.11%) | | |

**Supplementary Table S7: DESeq2 differentially expressed genes**

Chrom: chromosome; FC: log2 fold change; lfcSE: log2 fold change standard error; padj: Benjamini-Hochberg adjusted p-value

| **Gene Name** | **Chrom** | **FC** | **lfcSE** | **pvalue** | **padj** |
| --- | --- | --- | --- | --- | --- |
| *LRP2* | 2 | 2.7488846 | 0.37444169 | 3.00E-06 | 0.00426468 |
| *NKX2-2* | 20 | 6.04567979 | 0.8446254 | 2.32E-09 | 6.58E-06 |
| *P2RY1* | 3 | 2.92520385 | 0.31826531 | 1.46E-09 | 4.83E-06 |
| *POU3F4* | X | 3.98887224 | 0.63879036 | 2.88E-06 | 0.00426468 |
| *SALL1* | 16 | 4.4235896 | 0.75353452 | 5.54E-06 | 0.00647558 |
| *SIM1* | 6 | 4.35057431 | 0.73459814 | 5.09E-06 | 0.00632427 |
| *SLCO1A2* | 12 | 3.60823589 | 0.35000988 | 9.20E-14 | 4.57E-10 |
| *ZBTB16* | 11 | 2.42175637 | 0.35051568 | 4.99E-05 | 0.04508545 |
| *NOG* | 17 | 2.3699027 | 0.33437309 | 4.19E-05 | 0.03964386 |
| *NGEF* | 2 | 2.48465743 | 0.34796473 | 1.98E-05 | 0.0207636 |
| *PCDHACT* | 5 | 7.84882366 | 0.35969758 | 7.88E-81 | 1.57E-76 |
| *SERTAD4* | 1 | 2.68963284 | 0.31897171 | 1.18E-07 | 0.00029241 |
| *ZFHX4* | 8 | 2.74675703 | 0.33906257 | 2.58E-07 | 0.00053446 |
| *HAPLN3* | 15 | 3.03926447 | 0.45763783 | 8.35E-06 | 0.00922192 |
| *VSX2* | 14 | -2.9678774 | 0.38254876 | 2.69E-07 | 0.00053446 |
| *UNCX* | 7 | 4.63781157 | 0.56074403 | 8.73E-11 | 3.47E-07 |
| *KCTD4* | 13 | 4.77840219 | 0.81224789 | 3.29E-06 | 0.00436243 |
| *FAR2P1* | 2 | -2.2074591 | 0.13699566 | 1.21E-18 | 1.20E-14 |
| *LOC105371256* | 16 | 4.45924448 | 0.70640529 | 9.73E-07 | 0.00161278 |
| *LINC00237* | 20 | 6.25556025 | 1.07217595 | 9.50E-07 | 0.00161278 |
| *LOC107984886* | 16 | 6.57673776 | 1.31443486 | 2.21E-05 | 0.02195879 |
| *LOC112268271* | 20 | 4.63040918 | 0.45720547 | 2.01E-15 | 1.34E-11 |

**Supplementary Table S8: Summary of fusion genes involving *UBE3C* partial transcript**

Break: breakpoint; Split: split reads; Discord: discordant reads

| **Arriba** | | | | | | |
| --- | --- | --- | --- | --- | --- | --- |
| **Sample** | **Gene Name 1** | **Gene Name 2** | **Break 1** | **Break 2** | **Split** | **Discord** |
| P1 | UBE3C | LOC107986750(105280),  DPP6(118622) | 7:157187022 | 7:153629511 | 17 | 5 |
| P2 | UBE3C | LOC107986750(105280),  DPP6(118622) | 7:157187022 | 7:153629511 | 17 | 1 |
| P2 | UBE3C | LOC107986750(55056),  DPP6(168846) | 7:157187022 | 7:153579287 | 4 | 2 |
| P3 | UBE3C | LOC107986750(105280),  DPP6(118622) | 7:157187022 | 7:153629511 | 19 | 3 |
| P3 | UBE3C | LOC107986750(55056),  DPP6(168846) | 7:157187022 | 7:153579287 | 2 | 3 |
| **Defuse** | | | | | | |
| **Sample** | **Gene Name 1** | **Gene Name 2** | **Break 1** | **Break 2** | **Split** | **Discord** |
| P1 | UBE3C | PAXBP1P1 | 7:157187019 | 7:153629514 | 60 | 19 |
| P2 | UBE3C | PAXBP1P1 | 7:157187019 | 7:153629514 | 58 | 16 |
| P2 | UBE3C | PAXBP1P1 | 7:157188949 | 7:153579197 | 17 | 7 |
| P3 | UBE3C | PAXBP1P1 | 7:157187019 | 7:153629514 | 63 | 23 |
| P3 | UBE3C | PAXBP1P1 | 7:157188949 | 7:153579197 | 6 | 5 |
