## Supplementary Methods for "Novel gene-intergenic fusion involving ubiquitin E3 ligase UBE3C causes distal hereditary motor neuropathy: A new mechanism for motor neuron degeneration"

**Motor neuron progenitor induction and sMN differentiation**

iPSC colonies cultured to optimal density were dissociated using 0.5 mM EDTA, split 1:6 to 1:10 and seeded onto 6-well plates pre-treated with 0.167 mg/mL Matrigel. Cells were propagated for 3-4 days in TeSR-E8 iPSC culture medium until individual colonies reached optimal iPSC morphology. Optimal morphology is defined as colonies presenting with well-defined, uniform edges and comprising tightly compacted iPSCs with a large nucleus to cytoplasm ratio ^1-3^. Media and supplements used are summarised in Supplementary Table S3. Culture media was then replaced with a chemically defined neural base medium (NbM) comprising: 1:1 DMEM/F12 + Glutamax 1X (Gibco, Life Technologies) and Neurobasal medium (Gibco, Life Technologies), 0.5X N2 (Gibco, Life Technologies), 0.5X B27 (Gibco, Life Technologies), 0.1 mM L-ascorbic acid (Sigma) and 1X Penicillin/Streptomycin (Gibco, Life Technologies). NbM was supplemented with 2 µM Dorsomorphin (StemCell Technologies), 3 µM CHIR990021 (Sigma) and 2 µM SB431542 (StemCell Technologies). Supplemented NbM was changed the following day and then replaced every 48 h or as needed depending on the growth rate of the culture. Cells were maintained under these conditions in 5% CO_2_, humidified air at 37°C for up to 6 days. The cells were then dissociated using Dispase (5 U/mL, StemCell Technologies) diluted 1:5 in DPBS without Ca^2+^ and Mg^2+^ (DPBS-, Gibco, Life Technologies), split 1:5 to 1:6 and seeded onto Matrigel-coated 6-well plates. Cells were induced into MNPs by culturing in MNP medium comprising of NbM supplemented with 0.1 µM Retinoic Acid (RA, Sigma), 0.5 µM Smoothened Agonist (SAG, StemCell Technologies), 2 µM Dorsomorphin, 1 µM CHIR990021 and 2 µM SB431542 for 6 days in 5% CO2 humidified air at 37°C. MNP medium was changed the day after seeding and then replaced every 48 h for up to 6 days. At day 6, MNP were passaged 1:6 using Dispase (5 U/mL) diluted 1:5 in DPBS^-^ and expanded for 7 days in Expansion medium (ExM) comprising NbM supplemented with 0.5 mM Valporic Acid (VPA, Sigma), 0.1 µM RA, 0.5 µM SAG, 2 µM Dorsomorphin, 3 µM CHIR990021 and 2 µM SB431542. Alternatively, dissociated MNP were frozen in CryoStor CS10 at a split ratio of 1:2 to 1:3 to improve viability of cells post-thaw. After thawing, MNP were cultured in ExM and passaged once every 7 days for up to 5 passages.

To induce terminal differentiation of sMN from MNP, dissociated MNP were split 1:3 and seeded directly into ultra-low attachment 6-well plates (Corning-Costar) under suspension conditions and cultured in Neuronal Medium (NM) comprising of NbM supplemented with 0.5 µM RA and 0.1 µM SAG and cultured in suspension in ultra-low attachment 6-well plates (Corning-Costar). On the following day, suspensions cultures were further disaggregated to homogenise spheroid size by gentle pipetting and split 1:2 into ultra-low attachment 6-well plates. NM was replaced every 48 h thereafter until day 6 to produce homogeneous three-dimensional aggregates of neuronal cells (neuropsheres). After 6 days, neurospheres containing sMN were dissociated into single cells using dissociation solution (1:1 0.25% Trypsin-EDTA (ThermoFisher & Accumax (Thermofisher)) (ThermoFisher) and seeded onto Matrigel-coated plates. sMN were matured in Maturation Medium (MM) comprising of NbM supplemented with 0.5 µM RA, 0.1 µM SAG, 0.1 µM Compound-E (StemCell Technologies), 2ng/mL BDNF, 2ng/mL GDNF, and 2ng/mL CNTF (Life Technologies). After 72 h, MM was replaced and additionally supplemented with 0.01 µM SN38-P to purify cultures of proliferative progenitors and undifferentiated stem cells (Mao et al., 2018). sMN were cultured under these conditions for a further 4-9 days, replacing the media and supplementing with 0.02 µM SN38-P every 48 h.

**Nanostring nCounter gene expression assay.**

RNA samples (10 ng/µL) were mixed with target specific oligonucleotides, 3’-biotinylated capture probes and 5’-reporter probes tagged with a target-specific barcode and incubated for 16 h at 65°C to form target-specific ‘tag-RNA complexes’. The nCounter preparation station removed excess capture and reporter probes and the tag complexes were immobilised on a streptavidin-coated cartridge. The cartridge was scanned using the nCounter Digital Analyzer and the raw data was returned to our laboratory for analysis.

**Data visualisation.**

Nanostring and qPCR, immunofluorescence quantification and western blot quantification data was plotted using Graphpad Prism (v9). Data are expressed as mean normalised counts or as mean normalised counts ± standard error (SE) for biological replicates. For immunofluorescence quantification and western blot quantification, data was plotted using Graphpad Prism (v9) and expressed as the mean ± standard error (SE) for biological replicates. For mRNAseq studies, data was plotted in RStudio using a combination of DESeq2, EnhancedVolcano^4^, ggplot2^5^, gbeeswarm^6^ and pheatmap.^7^

**Additional *C.elegans* methods**

**Table 1:** List of *C. elegans* strains used in this study

| **Abbreviation** | **Strain** | **Genotype** | **Worm Base Acc.No.** |
| --- | --- | --- | --- |
| oxIs12 | EG1285 | oxIs12[unc-47p::GFP + lin-15(+)] |  |
| oxIs12; EmptyVector | MHB3 | nnaEx3[unc-25p::EMPTYBACKBONE]; oxIs12[unc-47p::GFP + lin-15(+)] | WBStrain00050609 |
| oxIs12; UBE3C-IF | MHB4 | nnaEx4[unc-25p::UBE3C-IF]; oxIs12[unc-47p::GFP + lin-15(+)] | WBStrain00050610 |

**Biochemical and locomotion behavior assays:**

Aldicarb and Levamisole paralysis assays were carried out as previously described.^8^ Day 1 adult animals were transferred to the 1 mM aldicarb and 0.2 mM levamisole assay plates and scored for paralysis every 10 min for a period of 160 min. To study animal locomotion, thrashing assay was carried out using day 4 old adults. Thrashing assay was carried out as previously described. ^8, 9^

**Heat stress assay:**

**Day 1 adult animals were used for heat stress assay. Sensitivity of the animals to heat stress was assayed by incubating the animals at 35ºC.^10^ The survival rate of the worms was assessed after 8 h. An animal was scored as dead by its unresponsiveness when prodded with a metal wire in the head and tail at least twice.**

**Live imaging and Neurodegeneration quantification assay:**

Age synchronised day 1 animals were anesthetised using 100 mM levamisole and mounted on a 3% agar pad. Leica DMI3000B inverted microscope and a ProgRes CF^Cool^ Camera were used for imaging. Scoring of the images for neurodegeneration was carried out as before.^8, 11^

**Statistical analysis:**

Graphpad Prism (v9) was used to carry out statistical analyses. One-way ANOVA and Tukey’s multiple correction test was used to calculate adjusted p-values unless mentioned otherwise. Statistical analysis of the neurodegeneration quantification was carried out as previously described.^12^ Corrected p-value < 0.05 was considered significant.
